# Spatial epitope barcoding reveals subclonal tumor patch behaviors

**DOI:** 10.1101/2021.06.29.449991

**Authors:** Xavier Rovira-Clavé, Alexandros P. Drainas, Sizun Jiang, Yunhao Bai, Maya Baron, Bokai Zhu, Maxim Markovic, Garry L. Coles, Michael C. Bassik, Julien Sage, Garry P. Nolan

## Abstract

Intratumoral variability is a seminal feature of human tumors contributing to tumor progression and response to treatment. Current technologies are unsuitable to accurately track phenotypes and subclonal evolution within tumors, especially in response to genetic manipulations. Here, we developed epitope combinatorial tags (EpicTags), which we coupled to multiplexed ion beam imaging (EpicMIBI) for *in situ* tracking of barcodes within tissue microenvironments. Using this platform, we dissected the spatial component of cell lineages and phenotypes in a xenograft model of small-cell lung cancer. We observed emergent properties from mixed clones leading to the preferential expansion of subclonal patches for both neuroendocrine and non-neuroendocrine cancer cell states in this model. In tumors harboring a fraction of PTEN-deficient cancer cells, we uncovered a non-autonomous increase of subclonal patch size in PTEN wildtype cancer cells. EpicMIBI can facilitate *in situ* interrogation of cell-intrinsic and cell-extrinsic processes involved in intratumoral heterogeneity.

## Introduction

Heterogeneity within a tumor can be result of genetic, epigenetic, and metabolic diversity, thus contributing to tumor progression and resistance to treatment (reviewed in (Easwaran, Tsai, and Baylin 2014; Jamal-Hanjani et al. 2015; Zahir et al. 2020)). For instance, multi-region sequencing of human tumors has revealed intricate patterns of tumor evolution: large proportions of region-specific genetic alterations were not detected in other areas of the tumors analyzed (Boutros et al. 2015; Gerlinger et al. 2012). The reversible process of epithelial-to-mesenchymal transition (EMT) provides another example of intra-tumoral heterogeneity. EMT is considered to be spatiotemporally regulated via epigenetic mechanisms, which in turn might influence cell migration ability and the initiation of metastasis (reviewed in (Derynck and Weinberg 2019)).

Tumors are also comprised of resident and infiltrating non-cancer cells, such as immune cells and fibroblasts (reviewed in (Hanahan and Weinberg 2011; Davidson et al. 2021)). This variable cellular organization, in conjunction with secreted biomolecules and extracellular matrix proteins, is defined as the tumor microenvironment (reviewed in (Ding et al. 2018; Hinshaw and Shevde 2019; Joyce 2005)). As interactions among different cell subtypes within the microenvironment will influence the pathobiology of cancer (Lei et al. 2020), defining the individual biologies of cancer subclones within the tumor microenvironment is an essential step towards providing context for the development of novel therapeutic strategies. The variability within tumors begs essential questions: do the subclones behave as lone actors, or is there an underlying, and interactive, order of these clones that aids tumor growth and survival? Is the tumor simply a competition amongst related peers, or is there a social ecology of tumor cells with important contributory, symbiotic, differences that must be present for success?

Single-cell technologies provide opportunities to investigate whether inherent order exists in tumor samples (reviewed in (González-Silva, Quevedo, and Varela 2020; Irish, Kotecha, and Nolan 2006)). Most single-cell sequencing technologies employ microfluidics for single-cell isolation (e.g., Fluidigm-based scRNAseq) or employ droplet-based barcoding of individual cells (e.g., Drop-seq) (Macosko et al. 2015). These technologies have revealed intra-tumor heterogeneity in cancer stem cells, EMT, immune load, clonal evolution, and response to treatment (González-Silva, Quevedo, and Varela 2020; Hong et al. 2019; Ireland et al. 2020). Since these cells are taken from their native context, such heterogeneity can appear random while the spatial organization was overlooked.

Recent advances have added in-depth spatial characterization of cellular subtypes within tissues by simultaneously measuring dozens of proteins or RNAs (Angelo et al. 2014; Eng et al. 2019; Rodriques et al. 2019; Goltsev et al. 2018; Schürch et al. 2020). Technologies such as Slide-seq quantify mRNA levels *in situ* (Rodriques et al. 2019) and hence distinguish between cell types; however, this approach cannot measure critical functional events such as post-translational modifications of proteins, or their translocation between cellular compartments. DBiT-seq is an alternative that can detect both mRNA and selected proteins (Liu et al. 2020), but it is currently limited to a 10-m scale that is unsuitable for subcellular compartmentalization.

Multiplex ion beam imaging (MIBI), can capture posttranslational modifications and measure over 40 proteins with a resolution down to 260 nm (Angelo et al. 2014; Keren et al. 2019). MIBI is thus suitable to assess active and inactive proteins within their subcellular locales, while simultaneously revealing phenotypic, epigenetic and metabolic cell states. Ideally, approaches such as MIBI can be combined with genetic modification of cell populations to allow perturbation analysis of individual cancer subclones to the behavior and organization of the tumor microenvironment. With enough scalability, combining multi-parameter acquisition and genetic screening would allow *in situ* detection of molecular drivers of cellular spatial identity, hence allowing large-scale testing of potential vulnerabilities in different regions of the tumor under investigation.

Lineage tracing methodologies have been useful for delineating mutations that occur during cancer evolution. CRISPR/Cas9 barcoding systems such as homing guide RNAs and CARLIN enable the reconstruction of a developmental lineage tree *in vivo* (Kalhor et al. 2018; Bowling et al. 2020; Quinn et al. 2021). The CRAINBOW system for fluorescently barcoding somatic mutations also provides a snapshot of oncogenic clonal expansion *in vivo*, suggesting that tumor heterogeneity is not random, but has some pre-determined characteristics that are heritable of reproducible (Boone et al. 2019). Other methods allow for visualization of the hypoxic state of cells, yielding cell-specific information about the local tumor cell states and the more global tumor microenvironment (Vermeer et al. 2020; Hartmann et al. 2021). However, such methods can be limited in the number of cells and parameters they can trace. Previous work have adapted protein barcodes for cancer cell clonality tracking via mass cytometry (Wroblewska et al. 2018), but is unable to resolve the spatial relationships of cells within a tissue.

Here, we report a method that couples an <u>epi</u>tope <u>c</u>ombinatorial <u>tag</u>ging strategy (EpicTag) with MIBI (EpicMIBI) for the spatial identification of cancer cell clones within a tumor. Cells overexpressing a unique combination of short epitopes are detected with MIBI using a robust multiplex antibody panel. EpicTags enable subclonal growth tracking and the visualization of a snapshot of tumor evolution. Using a xenograft model of small cell lung cancer (SCLC), an aggressive form of lung cancer with neuroendocrine features, we found that homogeneously tagged cells configure a tumor with reproducible phenotypically, epigenetically, and metabolically distinct cell types with certain phenotypes attaining dominance within tumor patches. Importantly, an emergent structure is apparent dependence of co-localized phenotypes that create an ecology with apparent co-residence of tumor subtypes. We also observed that loss a tumor suppressor, PTEN, can shift tumor evolution not only through cell intrinsic mechanisms but also by modifying behaviors of neighboring cancer cells that are PTEN wild type. These results underscore the necessity of imaging approaches such as EpicMIBI to characterize the ecosystem of the tumor microenvironment by delineating the individual contributions of cancer subclones towards global tumor burden, with clear opportunities for novel anti-cancer therapies.

## Results

### Imaging-based identification of specific cancer populations using epitope-based barcodes

To distinguish cell populations using epitope barcoding, we synthesized EpicTag vectors expressing a GFP reporter linked to a C-terminal tag consisting of combinations of three different epitopes (Figure 1A). We reasoned that coexpression of epitopes linked to GFP would ensure sufficient retention within cells to enable detection and cell identification by antibody-based multiplex imaging technologies. Each epitope combination tagged to GFP relates to a unique cell line through the corresponding combination of antibodies. We designed 20 different lentiviral vectors, each expressing three unique epitopes from a pool of six: AU1, FLAG, HA, StrepII, Prot C, and VSVg (Figure S1A). We then individually transduced human NCI-H82 SCLC cells with each construct, thus generating 20 EpicTag cell lines. Each of the 20 cell lines were then further barcoded with a unique combination of palladium isotopes, pooled at the same ratio, and labeled with a panel of isotope-conjugated antibodies targeting GFP and the six epitopes. Mass cytometry analysis showed GFP expression in the vast majority of the population (Figure S1B) and efficient detection of each of the six epitopes (Figure S1C), validating the robust expression from the vector backbone and the performance of the tag-specific antibodies. We applied UMAP (Uniform Manifold Approximation and Projection) (Becht et al. 2018) dimensional reduction to the epitope signals and detected groups of cells that express each of the 20 EpicTags (Figure S1D), indicating that all epitope combinations are expressed in the pooled population. As expected, each epitope-barcoded group contained a unique palladium barcode (Figure S1E, F). Taken together, these experiments validated the 20 EpicTag-barcoded NCI-H82 cell lines and the workflow necessary to generate and test barcoded cells.

**Figure 1.**
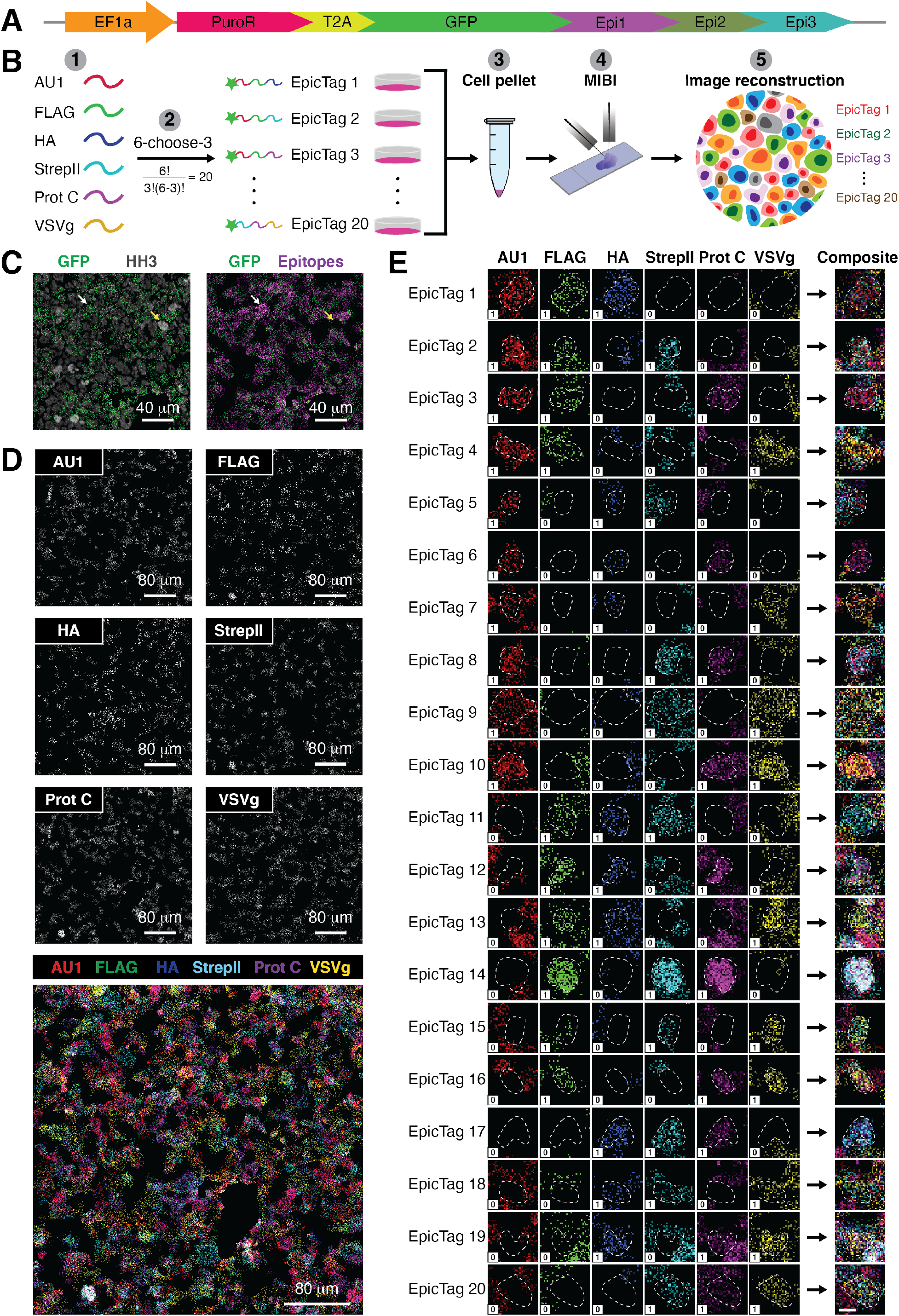
Imaging-based identification of specific cell populations with epitope-based barcodes. (**A**) Schematic of a representative EpicTag construct. The lentiviral-based vector contains an *EF1a* promoter (orange) for the expression of puromycin N-acetyl transferase (puroR; magenta), a T2A self-cleaving peptide (yellow), and GFP (green) fused to a C-terminal tag consisting of combinations of three short epitopes (epi 1, epi 2, and epi 3; purple, olive green, and cyan, respectively). (**B**) Workflow for multiplex imaging of epitope-based barcodes in cell pellets. (1) A library of short epitopes is selected. (2) Twenty cell lines expressing unique combinations of epitopes are generated based on a six-choose-three strategy. (3) Each cell line is grown *in vitro* in an independent flask, and then cultures are mixed and pelleted. (4) Cell pellet sections are mounted in gold-coated slides, and samples are stained with an isotope-conjugated antibody cocktail and loaded onto the MIBIscope. (5) A mass spectrum is obtained for each pixel of the targeted area, data are processed to obtain a two-dimensional image for each antibody, and epitope images are overlayed to identify barcodes in the image. (**C** to **E**) Representative MIBI images of a cell pellet consisting of 50% wild-type NCI-H82 cells and 50% of a pooled population of 20 barcoded NCI-H82 cell lines from the six epitopes depicted in Figure 1B.1. **C**) Left: Overlay of anti-GFP (green) and anti-Histone H3 (HH3) (gray) images showing cells with expression (white arrow) or absence (yellow arrow) of GFP. Right: Overlay of anti-GFP (green) and the sum of all six anti-epitope (magenta) images showing that epitope signals are detected in GFP-positive cells (white arrow) but not in GFP-negative cells (yellow arrow). Scale bars: 40 μm. **D**) Black and white images representative of anti-epitope (AU1, FLAG, HA, StrepII, Prot C and VSVg) images from the same field of view (FOV). The multicolor overlay at the bottom shows that each GFP-positive cell expresses a combination of epitopes. Scale bars: 80 μm. **E**) Multiple images showing a representative cell for each of the 20 barcodes. Each row is a barcode and each column is an epitope (AU1: red; FLAG: green; HA: blue; Strep II: cyan; Prot C: magenta; VSVg: yellow). The composite column is the overlay of the six anti-epitope images. Dashed lines were manually drawn to indicate the contour of the relevant cell. The numbers within the white squares indicate whether the epitope was expected (1) or not (0) in a given barcode. Images are enlargements from boxed regions in Supplementary Figure 1G. Scale bars: 10 μm.

Next, we used the 20 EpicTag-barcoded NCI-H82 cell lines and MIBI to distinguish barcoded cells in sections of barcoded cell pellets (Figure 1B). We first generated a cell pellet consisting of 50% wild-type NCI-H82 cells and 50% of a pooled population of the 20 EpicTag-barcoded NCI-H82 cell lines. MIBI analysis identified cells with and without GFP expression (Figure 1C, left), enabling identification of genetically modified cells within the pellet. The signal from the sum of all epitopes overlapped with the GFP signal and was absent from areas without GFP (Figure 1C, right), indicating that epitope signals are specifically found in GFP-containing cells only. We detected signals for each of the six epitopes, and the composite of all six revealed a complex multicolor image (Figure 1D). We identified all 20 barcodes within the cell pellet (Figure 1E and Figure S1G), validating EpicMIBI as a technique for imaging-based barcode detection. These *ex vivo* data indicate that combinatorial staining of epitopetagged proteins, coupled to a multiplexed imaging technology such as MIBI, enables the retention of spatial information for barcoded cells.

### Cells barcoded with EpicTags are detected *in vivo*

We determined the potential of the EpicTag strategy *in vivo* by pooling the 20 EpicTag-barcoded NCI-H82 cell lines to generate tumors in mice. Single-cell suspensions from these tumors were subjected to mass cytometry. All 20 combinations were detected in GFP-expressing cells (Figure S2A-D), indicating that EpicTag expression is durable in NCI-H82 cells within tumors *in vivo*. In tumor sections analyzed by MIBI, we detected all six epitopes, and cells expressed all the expected combinations of the three epitopes (Figure 2A, B). Thus, EpicMIBI can be leveraged as a proxy for common cell ancestry at the local scale amongst cancer cells in a proliferating tumor in mice.

**Figure 2.**
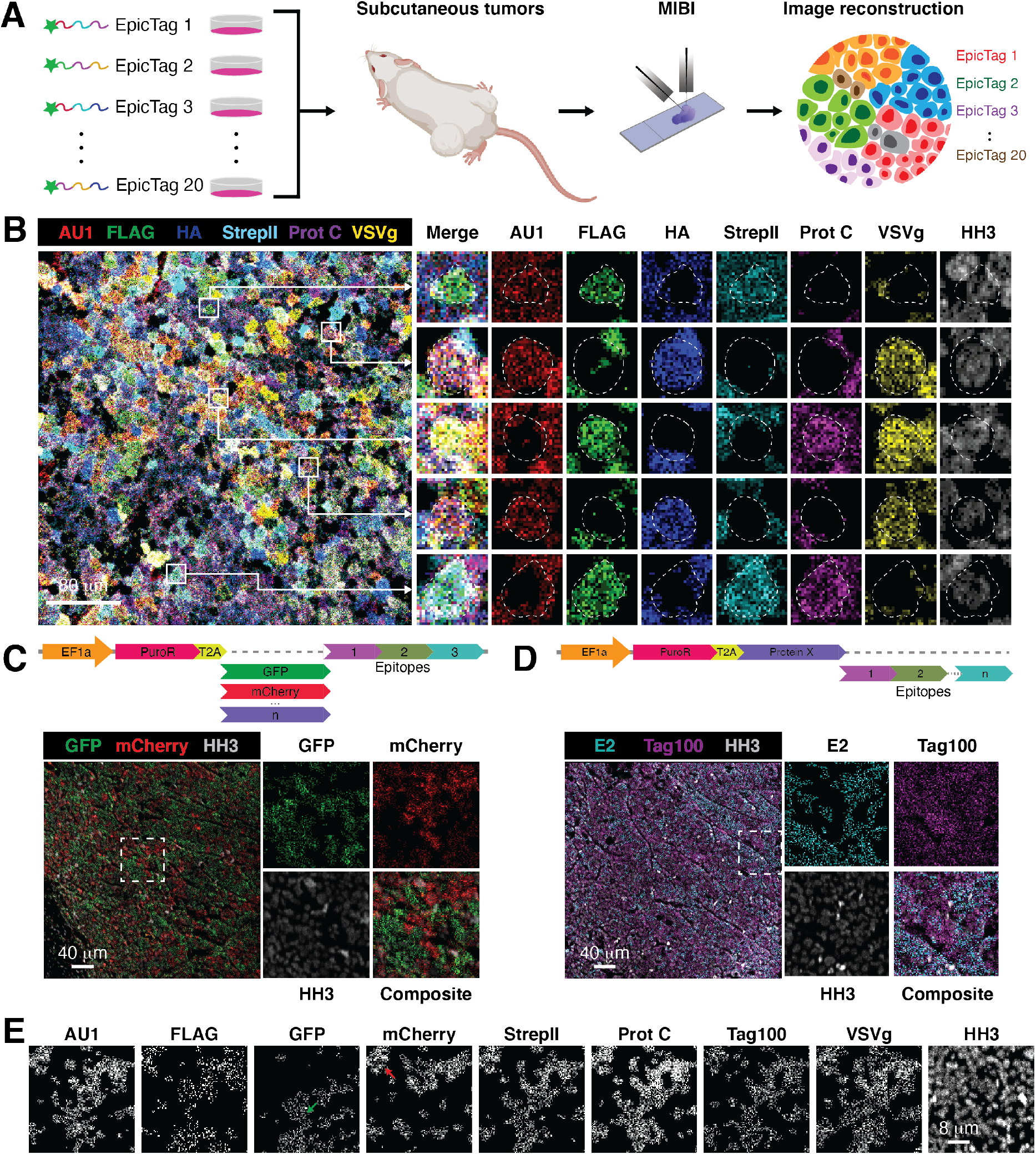
Detection of epitope-based barcodes in SCLC xenografts. (**A**) Workflow for multiplex imaging of epitope-based barcodes in subcutaneous NCI-H82 SCLC xenografts. (1) Cell lines expressing unique combinations of epitopes are generated based on an n-choose-k strategy and grown *in vitro*. (2) Barcoded cells are mixed and injected subcutaneously into the left and right flanks of a mouse. (3) Tumors are formalin-fixed, paraffin-embedded (FFPE), and sections are mounted in gold-coated slides. Samples are stained with an isotope-conjugated antibody cocktail and loaded onto the MIBIscope. (4) A mass spectrum is obtained for each pixel of the targeted area, data are processed to obtain a two-dimensional image for each antibody, and epitope images are overlayed to identify barcodes in the image. (**B**) Left: Representative overlay of AU1, FLAG, HA, StrepII, Prot C, and VSVg MIBI images of a tumor consisting of a pooled population of 20 barcoded NCI-H82. Right: Multiple enlarged images from boxed regions in the left image showing that individual cells express three of the six epitopes. Columns are overlay, with each of the epitopes (AU1: red; FLAG: green; HA: blue; Strep II: cyan; Prot C: magenta; VSVg: yellow), and HH3 (gray) to identify the cell nucleus. Scale bar: 80 μm. (**C-E**) Representative MIBI images of a tumor consisting of a pooled population of wild-type NCI-H82 cells and four barcoded NCI-H82 cell lines expressing GFP or mCherry tagged to six epitopes (AU1, FLAG, StrepII, Prot C, Tag100, and VSVg) or to three epitopes (FLAG, HA, and E2). **C**) Top: Schematic of a representative EpicTag construct. The vector contains an *EF1a* promoter (orange) for the expression of puromycin N-acetyl transferase (puroR; magenta), a T2A self-cleaving peptide (yellow), and a protein (GFP in green; mCherry in red; “n” number of proteins in violet) fused to a C-terminal tag consisting of combinations of three short epitopes (purple, olive green, and cyan). The dashed gray line indicates the modularity of the system. Bottom left: Overlay of anti-GFP (green), anti-mCherry (red), and anti-HH3 (gray) images showing cells expressing GFP or mCherry but not both simultaneously. Bottom right: Individual and enlarged images from boxed region on the left. Scale bar: 40 μm. **D**) Top: Schematic of a representative EpicTag construct with distinct epitopes. The vector contains an *EF1a* promoter (orange) for the expression of puroR (magenta), a T2A self-cleaving peptide (yellow), and a protein (violet) fused to a C-terminal tag consisting of combinations of “n” short epitopes (olive green, and cyan). The dashed gray line indicates the modularity of the system. Bottom left: Overlay of anti-E2 (cyan), anti-Tag100 (magenta), and anti-HH3 (gray) images showing cells expressing Tag100 or E2 but not both simultaneously. Bottom right: Individual and enlarged images from boxed region on the left. Scale bar: 40 μm. (**E**) Black and white images of the same field of view (FOV) showing cells expressing either GFP (green arrow) or mCherry (red arrow) and the six epitopes (AU1, FLAG, StrepII, Prot C, Tag100 and VSVg). Scale bar: 8 μm.

EpicTags are composed of two critical building blocks: a structured protein and a tail of epitopes. Expressing proteins alone as barcodes suffers from a lack of scalability, whereas short epitopes by themselves are not efficiently expressed or retained within the cell. This co-dependency can be leveraged to increase the number of barcodes based on experimental needs. On the protein side, distinct proteins can be tagged with combinations of three epitopes (Figure 2C, top). For example, we tagged mCherry with the six-choose-three (^6^C_3_) scheme described above (Figure S1A) and generated 20 additional EpicTag-barcoded NCI-H82 cell lines. We pooled the 20 mCherry^+^ and the 20 GFP^+^ cell lines and confirmed the presence of all 40 cell lines using mass cytometry (Figure S2E). We further validated the system by generating a pool excluding the NCI-H82 cell lines that were mCherry^+^ AU1-FLAG-HA^+^ and GFP^+^ StrepII-ProtC-VSVg^+^. We recovered the 38 expected combinations but not the two missing combinations (Figure S2F), highlighting the robustness of the system. We then generated tumors in mice by mixing mCherry^+^ and GFP^+^ cancer cells and identified mCherry-expressing and GFP-expressing cells (Figure 2C). These data show that the EpicTag system can be expanded to other proteins that can be tagged and detected using an antibody. We next expanded the number of short epitopes suitable for MIBI (Figure 2D, top). We generated vectors expressing E2 and Tag100 epitopes, generated tumors, and analyzed the samples. Both epitopes were detected *in vivo* (Figure 2D). We then generated tumors including cells expressing GFP or mCherry tagged with six epitopes (AU1, FLAG, StrepII, Prot C, Tag100 and VSVg). We detected cells expressing all six epitopes together and either GFP or mCherry (Figure 2E). Thus, a twelve-choose-six (^12^C_6_) scheme, or 924 combinations, is feasible given enough validated epitopes.

Altogether, these experiments demonstrate the modularity of the EpicTag system for tracing cancer-cell lineages within tumors through the diverse combination of well-expressed proteins tagged with a wide range of epitopes.

### Complex spatial tumor structures arise from a well-established cancer cell line

NCI-H82 cells belong to the NEUROD1-high subtype of SCLC (SCLC-N) (Rudin et al. 2019). Similar to other subtypes of SCLC, SCLC-N tumors are heterogeneous and harbor cancer cells with neuroendocrine (NE) and non-neuroendocrine (non-NE) features (Ireland et al. 2020; Stewart et al. 2020). Single-cell RT-qPCR analyses suggested that cells with NE and non-NE features co-exist in NCI-H82 cells in culture (Lim et al. 2017; Groves et al. 2021). Accumulating evidence indicates that the interplay between NE and non-NE cells is critical for tumor growth and evolution in SCLC (reviewed in (Shue, Lim, and Sage 2018)).

Using NCI-H82 xenografts as a model, we investigated the spatial organization of tumor heterogeneity using EpicMIBI, focusing on NE and non-NE phenotypes. To this end, we simultaneously stained barcoded NCI-H82 tumors with a 22-antibody MIBI panel to identify cell barcodes as well as various cell types in the tumor microenvironment and functional states for the cancer cells (e.g., cell cycle, differentiation) (Figure 3A). All the antibodies tested generated detectable signal with the expected cellular localization (Figure 3B). NCI-H82 cells were identified by GFP expression and further confirmed using a human-specific mitochondrial marker, which does not stain for mouse cells (Figure 3C). For each NCI-H82 cell identified, barcode expression (AU1, FLAG, HA, StrepII, Prot C, and VSVg) was analyzed, as well as markers of NE and non-NE differentiation (synaptophysin and vimentin, respectively), epigenetic states (H3K4me2, H3K27ac, and H4K8ac – histone modifications linked to active transcription), and metabolic states (GLUT1 and citrate synthase – markers for glycolysis and citric acid cycle activity, respectively) (Figure 3D). We also identified proliferative cells (Ki67 and the mitosis marker phospho-Ser28 of histone H3, HH3) and cells with DNA damage (phospho-Ser139 H2AX, γH2AX) (Figure 3E). In addition, we identified mouse endothelial cells (CD31) and stromal cells (aSMA) (Figure 3C). Overall, we analyzed 16 tiles, each one consisting of square field of views (FOVs) of 1200 μm to obtain the spatial location and phenotype of >200,000 cells in NCI-H82 xenografts (Figure S3A, B).

**Figure 3.**
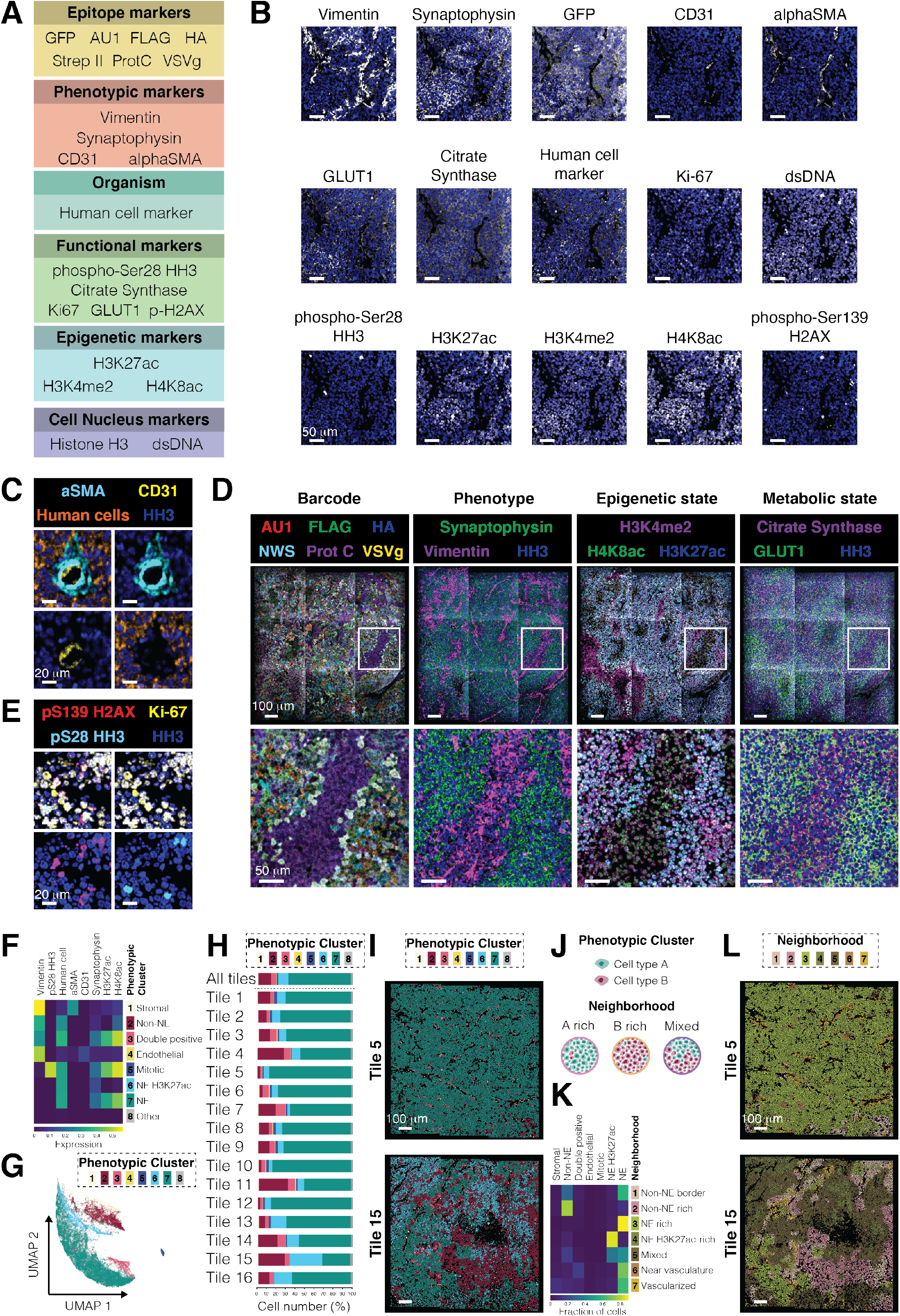
Spatial organization of NCI-H82 SCLC xenografts. (**A**) Summary of the antibody panel used for this study: Epitope markers for tracing cancer cell lineages; Phenotypic markers for distinguishing cell types and states; Organism markers to distinguish between human (NCI-H82 SCLC cells) and mouse cells; Functional markers for assessment of proliferation, DNA damage and metabolic state; Epigenetic markers for gene expression activity; Cell nucleus markers to stain DNA. (**B**) Images for each marker (gray) shown individually in a representative region of an NCI-H82 xenograft (Tile 3, field of view, FOV 2). HH3 is shown in all images (blue). Scale bars: 50 μm. (**C**) Images for aSMA (cyan), CD31 (yellow), and human cell marker (orange) in a representative region of an NCI-H82 xenograft (Tile 7, FOV 5). HH3 is shown in all images (blue). Scale bars: 20 μm. (**D**) Images for 14 markers in a representative region of the dataset (Tile 14). These data exemplify antibody staining patterns used to assess barcode (left), phenotype (second to the left), epigenetic state (second to the right), and metabolic state (right) of cells within an NCI-H82 xenograft. Scale bars: 100 μm (top) and 50 μm (bottom). (**E**) Representative images of pS139 H2AX (red; indicative of extensive DNA damage), Ki67 (yellow; indicative of proliferation), and pS28 HH3 (cyan; indicative of mitosis) in a representative region of the dataset (Tile 2, FOV 3). HH3 is shown in all images (blue). Scale bars: 20 μm. (**F**) A heatmap of mean marker expression for the eight phenotypic clusters (rows) and the eight markers that define them (columns). The intensity of each marker is defined by the color bar at the bottom (normalized counts). (**G**) UMAP of all cells in the dataset (n = 231715) colored by their phenotypic cluster. Each point represents a cell. Cells were grouped based on the expression of 13 markers (Vimentin, pS28 HH3, human cell marker, Ki67, aSMA, CD31, Citrate Synthase, GLUT1, pS139 H2AX, Synaptophysin, H3K27ac, H3K4me2, and H4K8ac). (**H**) Frequencies of each phenotypic cluster in the entire dataset (n = 231,715) and in each individual tile (Tiles 1 to 16). The dashed line separates the entire dataset representation from the individual tiles. Phenotypic clusters are indicated by color. (**I**) Representative phenotypic cluster maps on a region enriched in NE cells (Tile 5; phenotypic cluster 7) and a region with distinct cell types (Tile 15; phenotypic cluster 1 to 8). Scale bars: 100 μm. (**J**) Schematic of cellular neighborhoods. Cell type A (green) and cell type B (pink) might be in a region rich in homotypic (Neighborhood; A rich and B rich, respectively) or heterotypic (Neighborhood; Mixed) interactions. (**K**) A heatmap of the frequencies of each phenotypic cluster (columns) for each of the seven CNs (rows). The normalized intensity of each marker is defined by the color bar. Representative neighborhood maps on a region enriched in homotypic NE cell interactions (Tile 5; neighborhood 3) and in a region with distinctive substructures (Tile 15; neighborhoods 1 to 7). Scale bars: 100 μm.

Cell populations were defined using unsupervised self-organizing map (SOM) on the expression data from all tiles. We identified eight phenotypic clusters: seven SOM clusters based on marker expression, and a final manual annotation step to identify CD31^+^ cells (Figure 3F). Phenotypic clusters 2, 3, 5, 6, and 7 were of human origin, and clusters 1 and 4 were of mouse origin. Cluster 8, consisting of a 0.7% of the total cells, contained cells with low levels of expression of all markers, and it was therefore removed from the analysis. Cluster 1 contained mouse stromal cells as indicated by the expression of aSMA and vimentin, and lack of expression of the human-specific mitochondrial marker. Cells in cluster 4 were mouse endothelial cells as indicated by the expression of CD31 and vimentin, and the absence of the human-specific mitochondrial marker. Non-NE NCI-H82 cells (marked by vimentin and undetected expression of synaptophysin) were found in cluster 2, whereas NE cells (marked by synaptophysin and undetected expression of vimentin) were split into clusters 6 and 7. The split into two NE clusters was mainly driven by differences in the expression of epigenetic markers H3K27ac and H4K8ac, which were expressed in cluster 7 but not in cluster 6. Cluster 5 was composed of mitotic cells (marked by pS28 HH3). Cells in cluster 3 had high expression of both synaptophysin and vimentin. Visual inspection of cells in cluster 3 revealed that most of them correspond to NE cells near stromal cells, suggestive of a possible leakage of signal from stromal cells into NE cells, even though some of these cells may also represent a transition between the NE and non-NE states (Figure S3C). UMAP dimension reduction (Becht et al. 2018) was used to obtain a visual overview of where cells from each phenotypic SOM cluster were located. This representation shows that cells from each phenotypic SOM cluster are grouped together (Figure 3G), further supporting the clustering results.

Next, we calculated the frequencies of cells from each phenotypic SOM cluster in each tile analyzed (Figure 3H) and mapped the localization of the clusters onto each tile (Figure 3I and Figure S3D). Mouse stromal and endothelial cells (clusters 1 and 4, respectively) were present in all tiles analyzed (Figure 3H) and were scattered in the cluster-colored maps (Figure 3I). As expected, more than 75% of the cells in all tiles combined were NE cells (clusters 6 and 7) (Figure 3H, “All tiles”), and these cells were also the major populations in each individual tile (Figure 3H, “Tiles 1 to 16”). NE cells with high expression of H3K27ac (a marker of active transcriptional states) (cluster 7) were a major population in all tiles (Figure 3H, I), whereas NE cells with low levels of expression of H3K27ac (cluster 6), indicative of a distinct epigenetic state, were only present at a high frequency in a subset of tiles (Figure 3H, I; e.g., Tile 15). Similarly, non-NE cells (cluster 2) only presented at a high frequency in some tiles (Figure 3H, I; e.g., Tile 15). These results indicate that the distinct cancer cell states observed in the NCI-H82 xenografts are not homogenously distributed within the tumors. Thus, there may be specific local interactions between or within specific cancer cell states.

From an analytical standpoint, spatial cell arrangements might arise when two or more phenotypes are considered. In a reductionist view, if two phenotypes, A and B, are present in a tumor, they can be located in regions rich in A, rich in B, or containing both phenotypes (Figure 3J). Complexity escalates dramatically with increasing layered classifications or additional phenotypes. We previously showed that a tissue can be represented as a collection of cellular neighborhoods (CNs) (Schürch et al. 2020). CNs are regions of cells with a similar surrounding. The CN analytical framework defines distinct locations within tissues and enables a quantitative study of how neighborhoods influence function. Phenotypes identified in the NCI-H82 xenografts were clustered based on their 30 nearest neighbors to obtain seven distinctive CNs (Figure 3K) that we mapped to the segmentation map for each tile for a visual overview of the data (Figure 3L and Figure S3E). Certain CNs were enriched in cells with homotypic interactions: NE cells with high expression of H3K27ac (CN 3), NE cells with low levels of H3K27ac (CN 4), and non-NE cells (CN 2) (Figure 3K, L). Other CNs were composed of several cell types and states. For example, CN 1 included NE and non-NE cells (Figure 3K, L). CN 7, enriched in mouse cells, indicated vascularized and stromal regions of the tumor (Figure 3K, L). CN 6 was enriched in cells from cluster 3 (“double positive cells”), suggesting cells in this CN are surrounding the stroma (Figure 3K, L). CN 5 was composed of a mixture of NE cells with high and low expression of H3K27ac and non-NE cells was also identified (Figure 3K, L). Overall, a 52.9% of cells within NCI-H82 xenografts were in neighborhoods with homotypic interactions (CN 2, 3, and 4).

These analyses show that complex spatial rearrangements can arise *in vivo* in a tumor even from a well-established cancer cell line, raising the question of whether clonal expansion of cancer cells drive the formation of such structures.

### Clonal cancer cell growth is not spatially constrained in NCI-H82 xenografts

We reasoned that identifying the ancestries of the cells surrounding each cancer cell in the NCI-H82 xenograft model would provide insights into the processes that shape tumor architecture and growth. Barcodes in NCI-H82 follow a simple rule: three of the six epitopes are expected at a time in a cell. Advanced algorithms could be harnessed for cell debarcoding, but simply selecting a minimal image entity (e.g., a pixel, a segmented cell) and assigning the three most expressed epitopes provides an efficient and scalable pipeline for debarcoding.

By integrating cell- and pixel-based barcode assignment, we developed subclonal tumor maps to represent both the spatial locations of each cell within a tile, and its intrinsic EpicTag barcode information; the process is shown schematically in Figure 4A. In the cell-based barcode assignment single cells are first segmented based on histone H3 (HH3) and human-specific mitochondrial marker for nuclear and cytoplasm identification, respectively. Counts for each of the six epitopes within each cell are then sorted from highest to lowest. In the cell-agnostic pixel-based barcode assignment, sliding windows of distinct sizes are used to scan each pixel on the tile to identify regions enriched in barcodes. Subsequently, barcodes are assigned to individual pixels based on the three most highly expressed epitopes. Cell- and pixel-based barcode assignments are compared and merged following a set of rules (see methods section) to obtain a barcode for each cell within the tumor and a connectivity map for cells sharing a barcode and in physical contact. Using this pipeline, we identified all 20 possible epitope combinations on a tile from the NCI-H82 xenografts (Figure 4B, C and Figure S4A). Each barcoded population expressed the expected epitopes (Figure 4D), indicating that computational deconstruction of epitope-based images identifies common ancestries in the cancer cell population within the tumor.

**Figure 4.**
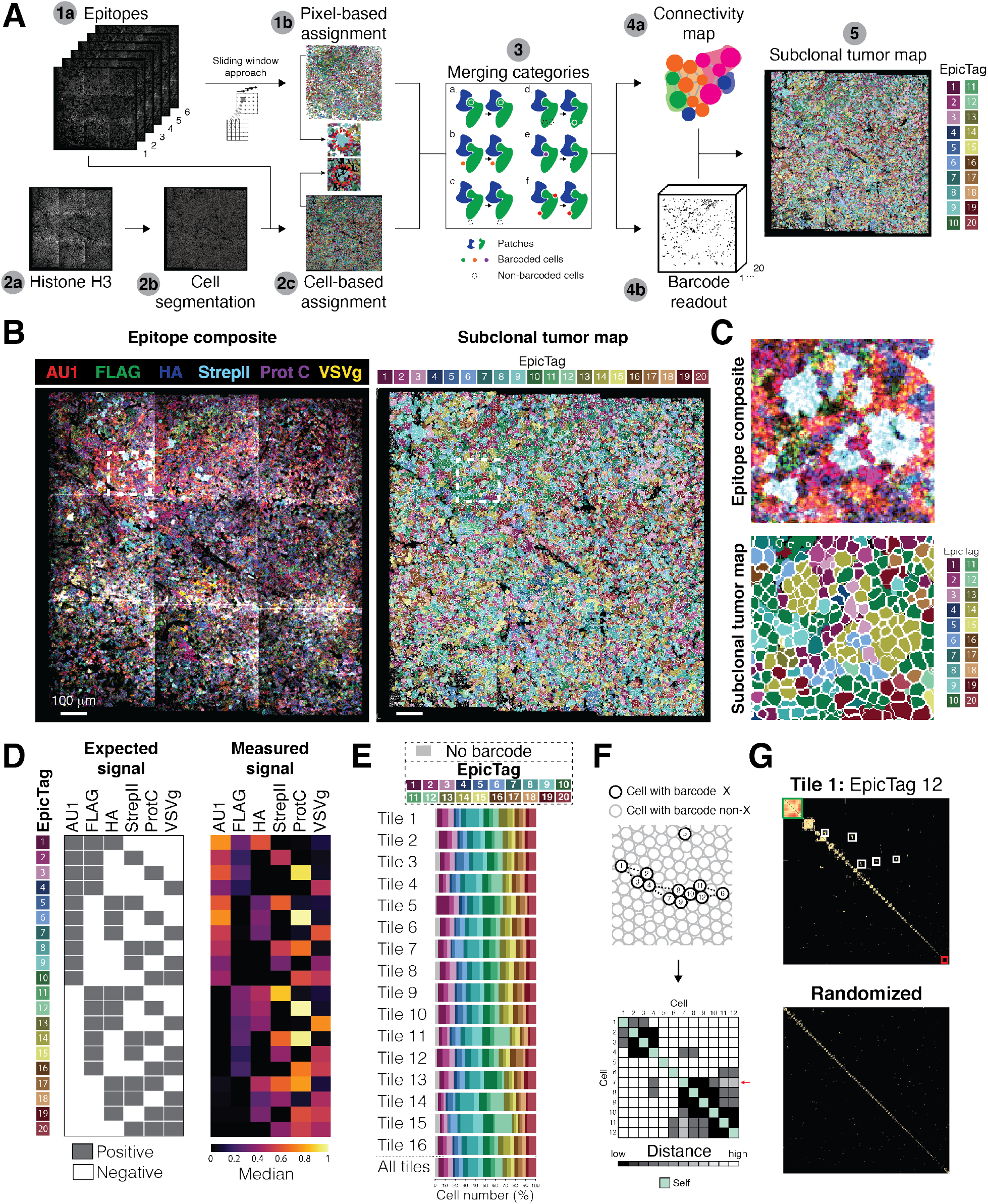
A debarcoding strategy identifies subclonal tumor patches. (**A**) Pipeline to reconstruct subclonal tumor maps. (1a) Epitope images are the input for pixel-based assignment of barcodes. (1b) A sliding window approach is applied to obtain a pixel-based assignment of barcodes. The cyan region within the dashed red circle in the enlarged image exemplifies a large region of the tumor sharing the same barcode. (2a) HH3 and human cell marker images are the initial inputs for cell-based assignment of barcodes. (2b) Single cells are segmented by the DeepCell neural network. (2c) Segmentation maps (step 2b) and epitope images (step 1a) are merged to obtain a cell-based assignment of barcodes. Epitope counts within each segmented region are extracted, normalized, and ordered to provide a barcode for each cell. Cyan cells within the dashed red circle in the enlarged image show that cells within the same patch (as shown in 1b) are not in direct contact after segmentation. (3) Merging step for cell- and pixel-based barcode assignments. Patches are shown as irregular shapes. Single cells are shown as small circles. Both depicted patches are bigger than a single cell to exemplify than patches consist of a group of cells. Barcodes are shown as colors. A dashed circle indicates a cell without an assigned barcode. To merge cell- and pixel-based barcode assignments, each cell within the tile is classified in one of six merging categories. On each category, left indicates the results from cell- and pixel-based barcode assignments and right indicates the result after integration. (4a) A connectivity map is obtained for each tile. Each circle represents a cell. Gray lines indicate that cells not directly touching in the segmentation map are from the same patch. (4b) A barcode is provided to each cell, resulting in a collection of images (one for each of the 20 barcodes). (5) Together, the connectivity map (4a) and the barcode readouts (4b) represent a subclonal tumor map. (**B-C**) Representative subclonal tumor map from a MIBI image of a tumor consisting of a pooled population of 20 barcoded NCI-H82 cell lines. **B**) Left: Overlay of anti-epitope (AU1, FLAG, HA, StrepII, Prot C, and VSVg) images showing barcode expression in the cells within the tumor. Right: Subclonal tumor map of the image on the left. Each cell was segmented and colored by barcode. Each EpicTag barcode has a distinct color (Barcode readout: 1 to 20). A white box indicates the area enlarged in Figure 5C. Scale bars: 100 μm. **C**) Top: Enlarged image of the epitope composite image in Figure 5B. The blueish colors identify regions enriched in FLAG, StrepII, and Prot C signals and missing AU1, HA, and VSVg signals. Bottom: Enlarged image of the subclonal tumor map in Figure 5B. The signals generating the blueish color in the epitope composite image on top are properly identified as EpicTag barcode 14 (olive green color). Segmentation provides a visual overview of the cells (white lines). (**D**) Quantification of epitope intensity on each debarcoded population from Figure 4B. Each row is an EpicTag barcode and each column is an epitope. Left: Expected signal for each epitope in each barcode. Grey indicates positive signal, and white indicates negative signal. For example, a population defined by EpicTag barcode 1 (first row) is expected to express AU1, FLAG, and HA. Right: Measured epitope signal in each debarcoded population. Color shows the median of the normalized counts. (**E**) Frequencies of each EpicTag barcode in the entire dataset (n = 231,715) and in each individual tile (Tiles 1 to 16). The dashed line separates the entire dataset representation from the individual tiles. The legend indicates the color for each EpicTag barcode. Top: Schematic representation of cells with the same EpicTag barcode. A black circle outlines a cell with a given barcode (“Cell with barcode X”). A grey circle outlines a cell without that barcode (“Cell with barcode non-X”). Each number indicates a cell. Cells with a particular barcode can be individually distributed (e.g., cell 1) or grouped (e.g., cells 2 to 4). Cells sharing a barcode can be distributed nearby (e.g., cell 6 and 11) or far apart (e.g., cell 5). Bottom: Grid of pairwise distances of the schematic shown on top. The numbers refer to the numbers of the schematic shown on top. A green box indicates self-interaction. A white box indicates the distance between two cells is higher than a threshold. The black to white gradient indicates distance (darker indicates closer). The red arrow highlights cell number seven. (**G**) Grid of pairwise interactions showing the distances of each cell to its fifth nearest neighbor for cells with EpicTag 12 in tile 1 (Top) and a randomized sample (Bottom). Randomization was performed by randomly assigning EpicTag 12 to segmented cells in tile 1, up to the number of EpicTag 12 in tile 1. Cells were arranged in the diagonal by patch size (larger patches in the top left corner). The green box exemplifies a subclonal tumor patch. The white boxes exemplify subclonal tumor patches that are closer in space. The red box exemplifies individually scattered cells.

**Figure 5.**
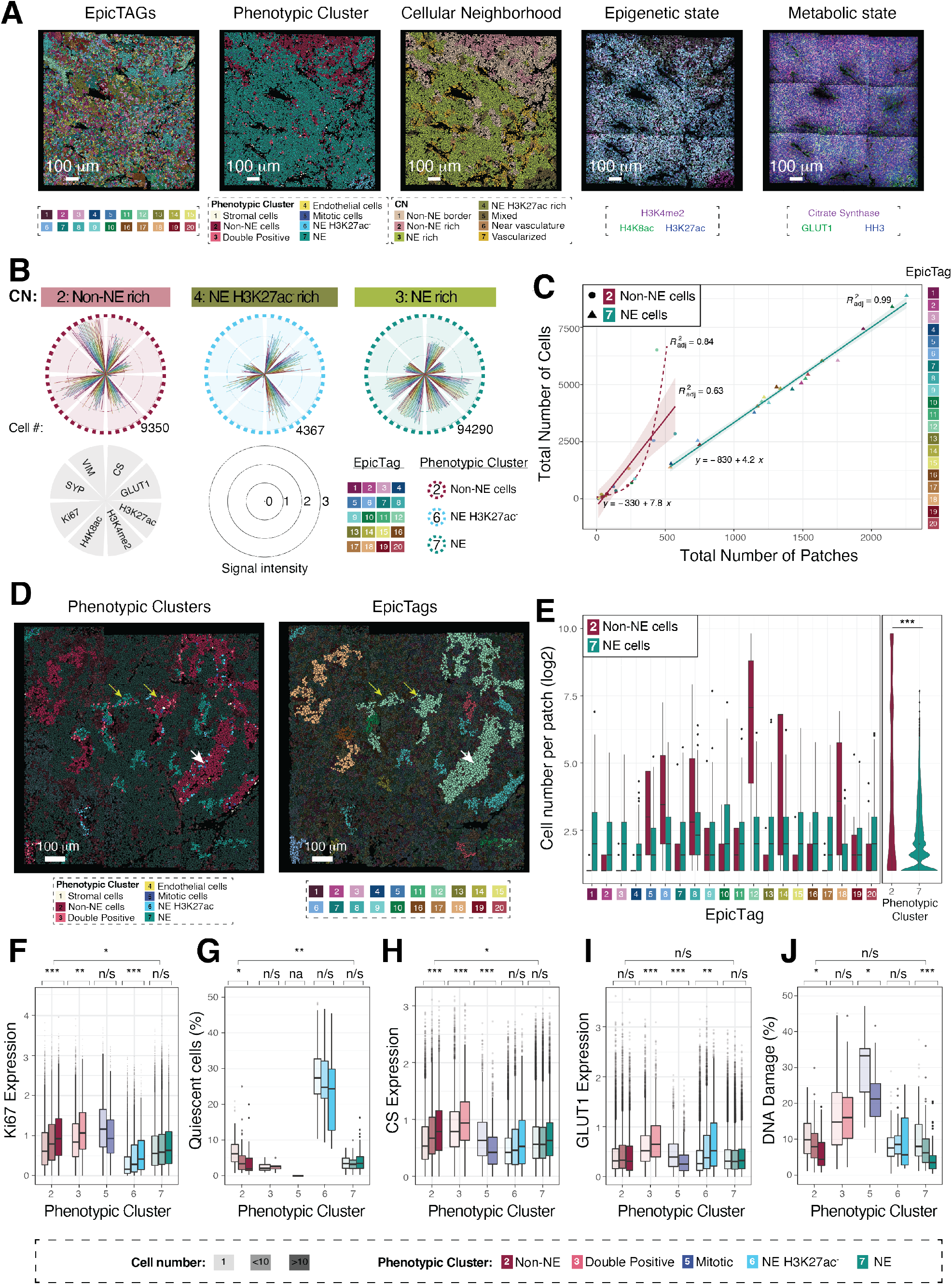
Non-neuroendocrine SCLC cells establish large patches with increased proliferative index and decreased DNA damage. (**A**) Overview of different classes of datasets. From left to right: Cell lineage assignment using EpicTag barcodes; phenotypic clusters; cellular neighborhoods (CNs); epigenetic markers, and metabolic markers. Scale bars: 100 μm. (**B**) Polar plots of marker expression (Vimentin (VIM), Synaptophysin (SYP), Ki67, H4K8ac, H3K4me2, H3K27ac, GLUT1, citrate synthase (CS)) in homotypic CNs (cluster 2 – CN 2, cluster 6 – CN 4, cluster 7 – CN 3). (**C**) The total number of barcoded cells (EpicTags 1 to 20) per patch from all tiles for clusters 2 (non-NE) and 7 (NE). Linear models fit of the data for NE and non-NE cells with R^2^ *adj* = 99% and 63%, respectively. An exponential fit was calculated for the non-NE cells yielding a fit of R^2^ *adj* = 84%. (**D**) Representative image with phenotypic clusters and EpicTag barcodes indicated. White arrow highlights a large patch of non-NE cells. Yellow arrows indicate where non-NE and NE cells share the same barcode. Scale bars: 100 μm. (**E**) Patch sizes by cell number per EpicTag for cluster 2 (non-NE cells, red) and cluster 7 (NE cells, cyan). p-value = 2.2*10^−16^. (**F**) Ki67 expression in individual clusters. For each cluster the data were separated by patch size: 1 cell, 1 to 10 cells, and over 10 cells. Significance is calculated by ANOVA within and between groups and adjusted by Bonferroni (Table S1). P-adj: 0.05-0.01:*, 0.01-0.001:**, <0.001:***. Quiescent cells per cluster. For each cluster the data were separated by patch size: 1 cell, 1 to 10 cells, and over 10 cells. Significance is calculated by ANOVA within and between groups and adjusted by Bonferroni (Table S1). P-adj: 0.05-0.01:*, 0.01-0.001:**, <0.001:***. (**H**) Citrate synthase expression. For each cluster the data were separated by patch size: 1 cell, 1 to 10 cells, and over 10 cells. Significance is calculated by ANOVA within and between groups and adjusted by Bonferroni (Table S1). P-adj: 0.05-0.01:*, 0.01-0.001:**, <0.001:***. (**I**) GLUT1 expression. For each cluster the data were separated by patch size: 1 cell, 1 to 10 cells, and over 10 cells. Significance is calculated by ANOVA within and between groups and adjusted by Bonferroni (Table S1). P-adj: 0.05-0.01:*, 0.01-0.001:**, <0.001:***. (**J**) Percentage of cells with DNA damage per cluster. For each cluster the data were separated by patch size: 1 cell, 1 to 10 cells, and over 10 cells. Significance is calculated by ANOVA within and between groups and adjusted by Bonferroni (Table S1). P-adj: 0.05-0.01:*, 0.01-0.001:**, <0.001:***.

We then processed the xenograft dataset with our debarcoding pipeline (Figure 4A) and obtained a subclonal tumor map for each tile (Figure S4B). After single-cell extraction and pooling of all tiles, we obtained the percentages of all twenty barcodes (Figure 4E). Cells with barcode 8 were the most highly represented, accounting for 9.71% of the pooled population. Those with barcode 4 were the least represented, accounting for 1.96% of the pooled population. The deviation from the expected 5% contribution from each of the 20 EpicTag-barcoded cell lines might be explained by several factors, for instance minor differences at initial seeding (Figure S4C), limited sampling, and presence of non-debarcoded events, which include non-cancer cells but also cancer cells without a clear barcode combination. All 20 EpicTag barcodes were identified on each of the tiles, but the distribution on a tile often deviated from the pooled data. For example, cells with barcode 1, which accounted for 5.52% of the pooled population, were 12.64% of the cells on tile 5 and 1.63% of the cells on tile 15 (Figure 4E). These data indicate that global tumor clonality is not a faithful representation of local clonal growth, thus suggesting that leveraging local scale as a variable may reveal novel mechanisms driving tumor evolution.

A subclonal tumor map conveys a static and fractional picture of the local complexity of cell-cell interactions within a tumor. Despite these limitations, certain local cancer cell behaviors that shape tumor growth can be inferred in such subclonal tumor map. Cancer cells sharing a barcode are expected to be detected both as individually distributed cells and as grouped in subclonal tumor patches of distinct sizes (Figure 4F, top). Further, individual cells and subclonal tumor patches with the same barcode are expected both in proximity to one another and scattered through the tumor (Figure 4F, top). This distribution is expected to arise in concert with the distribution from other cancer cells with distinct barcodes (Figure 4F, top; gray cells). Individually distributed cells represented 33% of cancer cells in the barcoded NCI-H82 xenografts (Figure S4D). The remaining 67% were found within subclonal tumor patches ranging in size from two cells to groups of hundreds of cells, with the absolute patch size rapidly declining (Figure S4D). We plotted the distance of each cell with barcode 12 to the fifth nearest neighbor with the same barcode in tile 1 and observed that most distances were smaller than the ones from a randomized distribution (Figure S4E). The same trend appeared when considering the nearest neighbors up to 300 (Figure S4F) and was shared among all barcodes in all tiles in the dataset (Figure S4G). Together, these analyses show that the introduced cancer cells are not randomly distributed in NCI-H82 xenografts and tend to form clonal patches.

Spatial cell distribution within a tumor can be viewed as a grid of pairwise distances (Figure 4F, bottom). This representation summarizes all existing distances below a given threshold, revealing clonal dynamics otherwise difficult to observe. For all cells in a barcode of a tile, we plotted the distances to the cells within the same patch and the distances to their nearest neighbors sharing a barcode. Some cancer cells sharing the same EpicTag grouped in relatively large, well-defined patches, whereas others were scattered individually around the tumor, far from cells of the same barcode. Certain patches co-existed near other patches of the same barcode (Figure 4G, top), and this spatial distribution did not occur in randomized data (Figure 4G, bottom). These results indicate that clonal growth is not spatially constrained, and further suggest that collective cancer cell movements within the tumor that may result from collective cell migration, growth of cells nearby, or other mechanisms, shape the tumor architecture.

### Non-neuroendocrine SCLC cells establish large clonal patches within the tumor

Simultaneous display of lineages (EpicTag barcodes), cell types and states (phenotypic SOM clusters), and CNs (neighborhoods), as well as subcellular visualization of functional markers, provide a rich dataset to interrogate drivers of tumoral evolution (Figure 5A). For an integrative data visualization of lineages, phenotypes, CNs, and marker expression, we plotted the expression of eight markers (vimentin, synaptophysin, Ki67, H4K8ac, H3K4me2, H3K27ac, GLUT1, and citrate synthase) for all barcoded cells from non-NE, NE H3K27ac^*low*^, and NE cell states (clusters 2, 6, and 7, respectively) in all CNs (Figure S5A). The coefficients of variation (CVs) of marker expression among barcodes in each cancer cell state and CN indicate that there is an overall similar expression of markers across cancer cell states and CNs with rare exceptions (Figure S5B, C), as would be expected for 20 isogenic cell lines.

We first focused on the three main cancer cell states (non-NE, NE, and NE H3K27ac^*low*^) in their respective homotypic CNs (non-NE rich, NE rich, and NE H3K27^*low*^ rich, respectively) (Figure 5B). There was comparable expression of most markers across all barcodes in the three selected clusters and CNs (Figure S5B, C), indicating good reproducibility between the EpicTag barcoded cells. A notable exception was high CV for H3K27ac for NE H3K27ac^*low*^ cells (cluster 6) in CN 4 (NE H3K27^*low*^ rich) (Figure S5C), but this marker is silenced in these cells and slight differences in expression might account for such a high CV. Outside the clusters in their homotypic CNs, CN 4 (NE H3K27^*low*^ rich) showed the highest CV (Figure S5B) (non-NE cells from cluster 2: 61%, and NE cells from cluster 7: 46%) indicating large marker expression variation within this CN. This might be explained by the low number of cells observed (Figure S5A; cell numbers for clusters 2 and 7 in CN 4 are 295 and 203, respectively). NE H3K27ac^*low*^ (cluster 6) and NE cells (cluster 7) in CN 2 (non-NE rich) also displayed increased CVs compared to the other CNs (Figure S5B; 60% and 23% for clusters 6 and 7, respectively), but this might also be explained by low numbers of cells (Figure S5A; cell numbers for cluster 6 and 7 in CN 2 are 380 and 895, respectively). Taken together, these analyses show good marker expression agreement amongst barcodes across clusters and CNs.

EpicTag-barcoded NCI-H82 cells generate patches of a broad range of sizes. We asked whether the total number of cells correlated with increased patch numbers and sizes for each EpicTag barcode and phenotypic cluster. In some cases, small patches might result from a tissue slice that captures cells only from the far edge of a larger phenotypic cluster. For NE cells (cluster 7), the number of cells and patches strongly correlated across barcoded cell lines (R*_adj_*^2^ = 99%) (Figure 5C and Figure S5D). In contrast, there was a lower correlation for non-NE cells (cluster 2; R*_adj_*^2^ = 63%) (Figure 5C and Figure S5D). An exponential model provided a better fit to the non-NE data (R*_adj_*^2^ = 84%) (Figure 5C), suggesting that the mechanisms of expansion differ for the NE and non-NE patches. The barcoded cells are from the same parent cell culture, and we expected similar barcode representations across the different clusters (with cells switching from one state to another similarly in all barcoded cell lines), which was observed for most phenotypic clusters (Figure S5E). One notable exception was the lack of correlation for the ratio between NE and non-NE cells (clusters 7 and 2, respectively) when comparing barcoded cell lines (Figure S5F), suggestive of specific dynamics in the acquisition of these cancer cell states within tumors.

The non-NE cells (cluster 2) generated fewer patches than the NE cells (cluster 7) (Figure 5C). The ratio of the slopes from the linear models indicates that for NE cells (cluster 7), 1.8 times more cells were in fewer patches than for non-NE cells (cluster 2), suggesting that non-NE cells are contained in larger patches (Figure 5D, white arrow). Indeed, the patch size was significantly larger in non-NE (cluster 2) compared to NE (cluster 7) and other cell states (Figure 5E, right, and Figure S5G, H). These data suggest that NCI-H82 cells with a non-NE phenotype remain together after cell division and tend to form larger patches.

We identified subclonal tumor patches with the same barcode in proximity that had both NE and non-NE states (clusters 7 and 2, respectively), suggesting a switch in phenotype (Figure 5D, yellow arrows). Non-NE cells (cluster 2) were identified in all barcoded populations (Figure 5 E), but six barcoded cell lines generated much larger patches of non-NE cells (cluster 2) compared to their NE counterparts (cluster 7) (Figure 5E, EpicTags 5, 6, 8, 12, 14, and 18) and four barcoded cell lines contained only 25-50 cells of non-NE cells (cluster 2) (Figure 5E, EpicTags 1, 3, 4, and 17), depicting a high variation in non-NE cell number across the 20 barcoded cell lines. Notably, approximately 90% of cells in each patch were assigned to the same cluster and the presence of a dominant cluster was significantly higher than expected based on a random distribution (Figure S5I). These observations suggest that NCI-H82 cells grown *in vivo* are mostly NE in nature but that large patches of non-NE cells may sometimes “stabilize” and expand in response to unknown stimuli.

We surmised that the generation of these large patches of non-NE cells (cluster 2) may be due to increased proliferation and thus examined markers associated with cell proliferation in tumor sections. Analysis of Ki67, a marker of non-quiescent cells (Miller et al. 2018), showed that mitotic cells (cluster 5) had the highest level of Ki67 expression as expected (Figure 5F, G). In non-NE cells (cluster 2), Ki67 levels significantly increased, and the number of quiescent cells significantly decreased with patch size (Figure 5F, G). In contrast, there was no significant correlation between Ki67 expression, number of quiescent cells and patch size for NE cells (cluster 7) (Figure 5F, G). NE H3K27ac^*low*^ cells (cluster 6) had a low proliferative index (Figure 5F, G), highlighting a functional difference with NE cells (cluster 7). Interestingly, non-NE (cluster 2) and NE cells (cluster 7) were more quiescent in CN 4 (18% and 11%, respectively), which is enriched in NE-H3K27ac^*low*^ cells (cluster 6) (Figure S5J, K). This suggests that the proliferation of NE and non-NE cells is affected by the presence of more slowly cycling NE-H3K27ac^*low*^ cells. Ki67 expression was higher in the larger patches across all CNs, although this was observed to a less extent for CN 4, enriched in NE-H3K27ac^*low*^ cells (cluster 6) (Figure S5L). Notably, citrate synthase (CS) expression was significantly elevated in larger non-NE (cluster 2) patches (more than 10 cells), but GLUT1 levels were similar across patch sizes in both non-NE (cluster 2) and NE cells (cluster 7) (Figure 5H and 5I), suggesting elevated activity of the citric acid cycle compared to glycolysis in larger non-NE patches. Finally, we noted that that larger patches (more than 10 cells) of non-NE (cluster 2) and NE cells (cluster 7) showed less DNA damage based on γH2AX signal (Figure 5K) independent of the neighborhood in which these cells were located (Figure S5M). Decreased DNA damage, changes in metabolism, and increased proliferation may all contribute to the expansion of larger non-NE subclonal patches once small patches have formed.

### PTEN loss promotes a non-cell autonomous increase of patch size in neighboring PTEN wild-type cancer cells

Our method for *in situ* lineage tracing (barcodes) coupled to phenotypic cell characterization and the accompanying pipeline for subclonal tumor patch analysis provide a unique opportunity to study functional interactions between genetically distinct cancer cells within the tumor microenvironment. PTEN is as a known potent tumor suppressor in SCLC (McFadden et al. 2014; Cui et al. 2014) but the evolution of PTEN mutant clones within SCLC tumors has not been examined. We knocked out *PTEN* by Cas9-sgRNA ribonucleoprotein delivery (*PTEN^−/−^*) in NCI-H82 SCLC cells expressing EpicTags 1 and 20 (Figure 6A) and generated six control cell lines by delivering non-targeting Cas9-sgRNA ribonucleoproteins (EpicTags 2, 4, 14, 16, 17, and 18). These eight cell lines were then pooled with the remaining twelve wild-type EpicTag NCI-H82 lines to generate a population consisting of 20 cell lines with individual expected contributions of 5%. A pool of the 20 original EpicTag lines was used as a control. In the pool containing the *PTEN^−/−^* lines, EpicTag lines 1 and 20 grew more rapidly in culture than the other lines (Figure 6B, C) and accounted at the end of the experiment for 15% and 29% of the population, respectively. The original EpicTags lines 1 and 20 did not have a competitive advantage (Figure 6C, left). These observations validated the expected tumor suppressor role for PTEN in SCLC cells.

**Figure 6.**
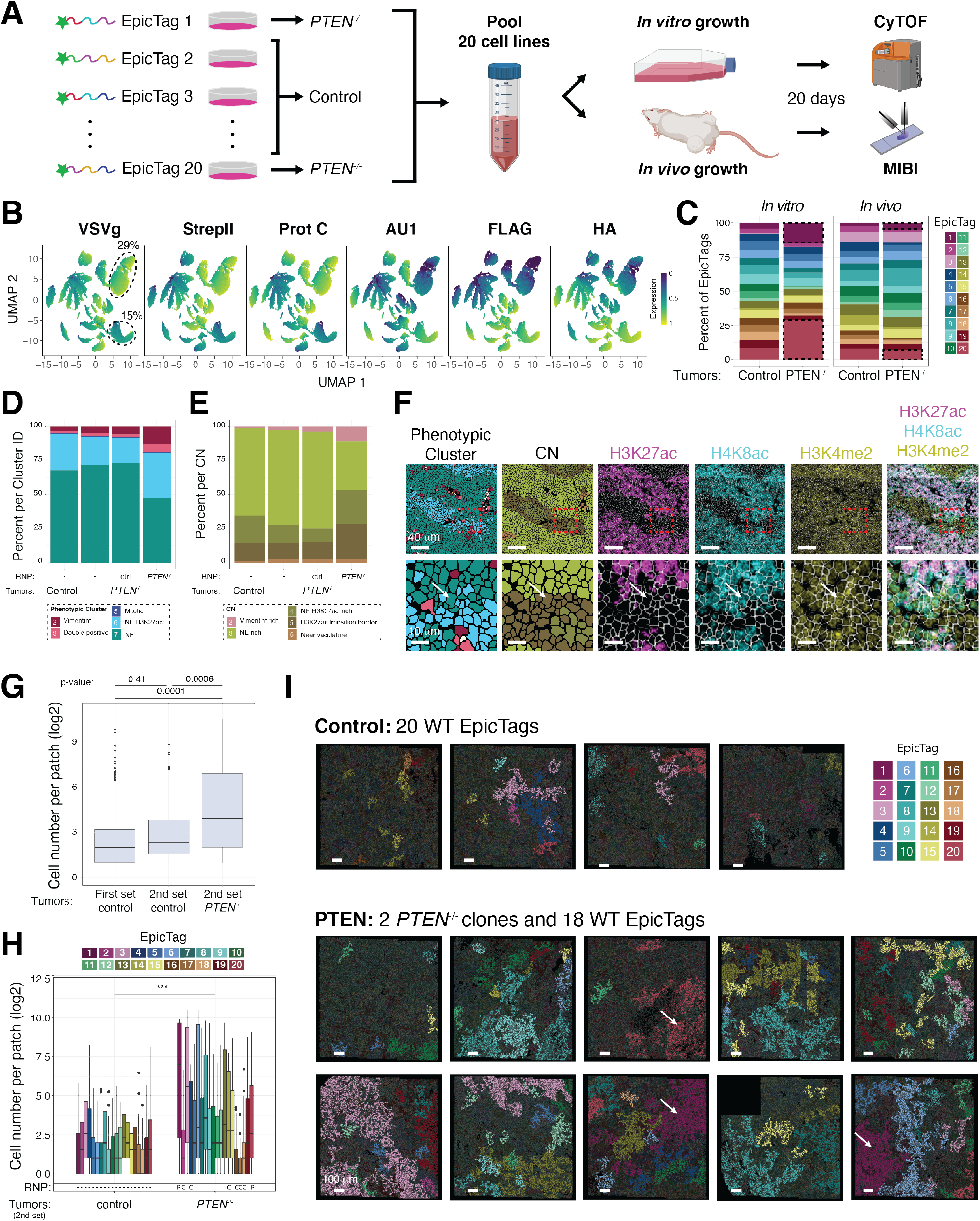
PTEN deficiency in a fraction of cancer cells modifies tumor architecture through cell intrinsic and extrinsic mechanisms. (**A**) Workflow for multiplex imaging of genetically modified epitope-based barcoded cells in subcutaneous NCI-H82 xenografts. (1) Cell lines expressing unique combinations of epitopes are generated based on an *n*-choose-*k* strategy and grown *in vitro*. *PTEN* was knocked out by RNP sgRNA:Cas9 nucleofection in cells expressing EpicTags 1 and 20. (2) Barcoded cells are mixed. (3) The same mixture is divided and grown *in vitro* or injected subcutaneously into the flanks of a mouse. (4) The *in vitro* culture is analyzed by mass cytometry. (5) Tumors are analyzed by MIBI. (**B**) UMAP of cultured cells (n = 69,065) grouped by epitope expression and colored by epitope (AU1, FLAG, HA, StrepII, Prot C, and VSVg). Dashed ellipses and percentages indicate EpicTag 1 (bottom) and 20 (top) and their respective contributions to the final population. (**C**) Frequencies of each EpicTag barcode in cells grown *in vitro* and *in vivo* in the “Control” consisting of 20 wild-type EpicTag barcoded cells and in the “*PTEN^−/−^*” pool consisting of *PTEN* knockout cells (EpicTags 1 and 20), unedited but nucleofected cells (EpicTags 2, 4, 14, 16, 17, and 18), and wild-type cells (EpicTags 3, 5, 6, 7, 8, 9, 10, 11, 12, 13, 15, and 19). The legend indicates the color for each EpicTag barcode. Black dashed boxes indicate *PTEN* knockout cells. (**D**) Frequencies of each phenotypic cluster in “Control” (n = 56,199) and “*PTEN^−/−^*” (n = 142,056) tumors. For the “*PTEN^−/−^*” tumors, frequencies for wild-type cells (“−”), unedited but nucleofected cells (“ctrl”), and *PTEN* knockout cells (“*PTEN^−/−^*”) are shown. (**E**) Frequencies of each CN in “Control” (n = 56,199) and “*PTEN^−/−^*” (n =142,056) tumors. For the “*PTEN^−/−^*” tumors, frequencies of wild-type cells (“−”), unedited but nucleofected cells (“ctrl”), and *PTEN* knockout cells (“PTEN*−/−*”) are shown. The legend indicates the color for each CN. (**F**) Phenotypic cluster map, CN map, and images of H3K27ac (magenta), H4K8ac (cyan), and H3K4me2 (yellow) in a representative region of a *PTEN^−/−^* NCI-H82 xenograft (Tile 4, FOV 4). Bottom row images are enlarged representations of red dashed squares in the top row. White arrows indicate representative H3K27ac^−^, H4K8ac^+^, and H3K4me2^+^ cells enriched in CN 5. Scale bars: 40 μm (top) and 10 μm (bottom). (**G**) Overall patch size by cell number in “Control” and “*PTEN^−/−^*” tumors. The overall patch size from the dataset shown in Figure 5 is shown as reference (“First set control”). P-values calculated by the student’s t-test. (**H**) Patch size by cell number per EpicTag in “Control” and “*PTEN^−/−^*” tumors. For “*PTEN^−/−^*” tumors, sizes of wild-type cells (“−”), unedited but nucleofected cells (“C”), and *PTEN* knockout cells (“P”) are indicated. The legend indicates the color for each EpicTag barcode. P-value calculated by the student’s t-test. (**I**) Images of subclonal tumor map with EpicTag barcodes highlighted in patches of more than 50 cells in “Control” and “*PTEN^−/−^*” tumors. White arrows indicate *PTEN* knockout patches. The legend indicates the color for each EpicTag barcode. Scale bars: 100 μm.

The pooled population containing the two *PTEN^−/−^* lines and the control pool were injected subcutaneously into mice to generate tumors. We analyzed approximately 198,000 cells from ten tumor regions that included *PTEN^−/−^* cells and four tumor regions with only control cells. All 20 barcodes were detected in tumors from both groups (Figure S6A). Surprisingly, in this context, EpicTag lines 1 and 20 did not expand more than control cells, and each EpicTag line accounted for a similar percentage in all tumors (Figure 6C, right). We also found no difference in the proliferation of *PTEN^−/−^* and control cells in the xenografts (Figure S6B). These data show that *PTEN* knockout in NCI-H82 cells provides a competitive growth advantage in culture but, interestingly, not *in vivo* in the context studied.

Analyses of 2D cultures can reveal causal links between genotype to function but may not recapitulate relevant characteristics of 3D *in vivo* tumor growth (Han et al. 2020). The relative lack of expansion of the *PTEN^−/−^* cells compared to the control cells in the xenograft models suggested that wild-type and PTEN^*−/−*^ cells may interact differently in 3D tumors than 2D cultures. However, the overall tumor architecture, as defined by SOM clusters and CNs, was comparable between tumors that included *PTEN^−/−^* cells and tumors that included only wild-type cells (Figure S6C–G), suggesting that no major differences in the spatial arrangements of the NCI-H82 xenografts were induced when 10% of the population lacked PTEN.

By barcoding cells, fine details of tumor architecture can be assessed. We analyzed patches in tumors that developed from pools containing *PTEN^−/−^* cells and controls (*PTEN* wild-type) and from tumors from pools of *PTEN* wild-type cells. Similar to our previous analysis (Figure S5I), we observed patches of cells with a dominant phenotype for the control (*PTEN* wild-type) cells (Figure S6H). In contrast, the patches of cells that lacked *PTEN* were generally composed of multiple cell states (Figure S6H), including patches with a mixture of NE (cluster 7), vimentin-positive (cluster 2), and NE H3K27ac^*low*^ (cluster 6) cells (Figure S6I). Notably, *PTEN^−/−^* cells had a higher percentage of NE H3K27ac^*low*^ cells (cluster 6) compared to control cells in the same tumor (Figure 6D). NE H3K27ac^*low*^ cells (cluster 6) were smaller than conventional NE cells (cluster 7) (Figure S6J) and were identified in areas with features of necrosis (Figure S6K). *PTEN^−/−^* cells were more likely to be located in CN 5 (Figure 6E), areas of the tumor corresponding to the border between regions rich in conventional NE cells (CN 3) and NE H3K27ac^*low*^ (CN 4) (Figure 6F). Cells in CN 5 expressed considerably less H3K27ac compared to cells in CN 3, and levels of H4K8ac were lower than in CN 3 and CN 4, although the difference was less dramatic (Figure S6L), suggesting a stepwise process of epigenetic alterations. Overall, these data suggest that loss of PTEN in cell populations *in vivo* is accompanied by both an increase in plasticity (or a lack of commitment to certain fates) and a propensity for higher cell death, which may account for the lack of observed competitive advantage of *PTEN^−/−^* cells *in vivo*.

Our patch analysis of control NCI-H82 xenografts (Figure 4) provided a ground truth for the expected subclonal growth dynamics of this cell line *in vivo* (Figure 6G; “First set control”). The same patch size was also observed in the control tumors generated in parallel to the tumors containing *PTEN^−/−^* populations (Figure 6G; “2nd set control”). Strikingly, the clonal patches were larger in tumors containing *PTEN^−/−^* cells than in tumors with only control cell lines (Figure 6G; “2nd set PTEN”). Not only were patches containing *PTEN^−/−^* cells larger, but there was also an increase of patch size for most *PTEN* wild-type control cell lines (Figure 6H, I). Larger patch size in control cells in tumors containing *PTEN^−/−^* cells could not be explained by increased proliferation (Figure S6M), altered GLUT1 or citrate synthase levels (Figure S6M), or higher amounts of DNA damage (Figure S6N). While the reason for the larger patch size remains unknown, and was unexpected, these data demonstrate that the overall architecture of NCI-H82 xenografts is altered by the presence of a *PTEN^−/−^* population initially seeded at ~10%. Modified tumor architecture is not just explained from a *PTEN^−/−^* cell intrinsic perspective: these genetically edited cells exert non-autonomous effects on other cancer cells within the tumor.

## Discussion

Clonal evolution is a major determinant of tumorigenesis. Increasing evidence suggests that the local microenvironment strongly influences clonal cancer cell growth, and the field will greatly benefit from strategies to track clonality *in situ* while keeping tissue context information. By applying a strategy for *in situ* tracking of barcodes in a xenograft model of SCLC, we show that lineage tracing integrated with deep *in situ* tissue characterization can reveal new insights into tumor architecture and mechanisms of tumor evolution. Our data show that even a well-established cell line grown in culture for decades can form complex tumor structures with apparently inter-dependencies wherein one or more phenotypic states prefer or generate nearby distinct phenotypic states. Within these structures, we identified localized growth of subclonal patches and detected distinct spatial dynamics of the identified cell phenotypes within the tumor. We further extended the capabilities of the system by linking barcode expression to specific genome edits, opening new opportunities to study, *in situ*, complex autonomous and non-autonomous phenotypes during tumor growth.

The NCI-H82 xenografts analyzed in this study had heterogeneity demarcated within distinct regions of the tumor. From a global non-spatially defined perspective, this heterogeneity is in line with previous studies indicating that NCI-H82 cells exist in a rather undefined region of the low-dimensional space from the continuum of functional states of several SCLC cell lines (Groves et al. 2021). In our dataset, we identified three major states of NCI-H82 cells: NE cells, NE cells lacking H3K27ac, and non-NE cells. NE cells were identified by synaptophysin expression and were further divided into cells that did and did not express H3K27ac. NE H3K27ac positive cells were present in all analyzed regions, whereas NE H3K27ac negative cells were enriched in certain regions of the tumors. The spatial component of NE H3K27ac negative cells, together with the non-clonal nature of these cells, suggests a coordinated H3K27ac shutdown directed by the microenvironment. In regions enriched in NE cells with high levels of H3K27ac, we observed a localized growth of subclonal patches, as well as grouping of individually barcoded cells, which is suggestive of spatial dynamics and phenotypic dependencies that remain to be understood. Cancer cells in the tumor core can be notably dynamic (Staneva et al. 2019). The ability to study how these movements relate to cancer cell clonal growth will have profound implications in our understanding of tumor evolution.

Non-NE NCI-H82 cells were identified by high levels of vimentin, a prominent mesenchymal marker (reviewed in (Dongre and Weinberg 2019)), suggesting that these cancer cells have undergone an epithelial-to-mesenchymal-like transition. Subclonal patches of non-NE cells were significantly larger than NE cell-containing patches. The non-NE cells in larger patches had an increased proliferation and decreased percentage of quiescent cells compared to non-NE cells in smaller patches or individually scattered cells, providing an explanation of how non-NE cells stabilize within the xenografts. Notably, non-NE cells in larger patches had increased citric acid cycle activity indicating that these cells may exploit the respiratory chain for energy production. Larger patches contained proportionally fewer cells with DNA damage cells than smaller patches or individual non-NE and NE cells, and hence cell fitness (as defined by low DNA damage) may lead to larger patches. Other factors may also play a role, such as preferred cell type targeting by the innate immune system. Notch signaling activation is related to the NE to non-NE shift, and this may be another mechanism in play (Lim et al. 2017).

PTEN is a prominent tumor suppressor in SCLC (McFadden et al. 2014; Cui et al. 2014). In NCI-H82 cells, we found that PTEN loss results in rapid expansion of SCLC cells *in vitro* but not *in vivo*. Notably, both *PTEN^−/−^* and wild-type cells formed large clonal patches in tumors containing *PTEN^−/−^* cells. The analysis suggests that increased patch size is independent of proliferation, DNA damage, GLUT1 and citrate synthase expression, which may hint that PTEN deficiency results in the secretion of paracrine factors that enhance the clonal growth of cancer cells in the proximity of *PTEN* mutant cells. Alternatively, PTEN mutant cells might create conditions that minimize mobility of cell clones, and thus reduce ‘fragmentation’ and mixing. Hence, our data support the idea that PTEN loss may alter cancer behaviors *via* cell-intrinsic and cell non-autonomous effects in the tumor microenvironment (Toso et al. 2014; Trimboli et al. 2009; Bronisz et al. 2011). Whether these phenotypes related to mesenchymal transitions and non-cell autonomous roles for PTEN specific to a few SCLC models or broadly relevant to SCLC remains to be determined.

The EpicMIBI approach is constrained by the number of barcodes deployed, the size of the antibody panel, constraints on the acquisition area, and debarcoding fidelity. We show *in situ* detection of twenty barcodes using GFP tagged to three out of six epitopes and provide evidence that this strategy could be easily extended *in vivo* to 140 barcodes through use of three additional antibodies. Our results also show codetection of six epitopes in the same tumor cell, suggesting that thousands of barcodes could be created by leveraging additional tags (e.g., T7, Universal, V5, HSV). Additionally, an increase in barcoded cell numbers can theoretically be obtained by controlling subcellular location (e.g., nucleus, membrane) (Loulier et al. 2014) or expression levels of the same barcode (Sakaguchi, Leiwe, and Imai 2018). Epitope sequences are currently introduced *in vitro* as unique and heritable barcodes, but alternatives similar to the Confetti reporter system (Snippert et al. 2010) would allow *in vivo* expression of thousands of stochastic barcodes in a temporally and spatially controllable manner.

We leveraged the multiplex imaging capabilities of MIBI to detect EpicTags as well as a panel of 22 antibodies covering a range of biological functions (e.g., epigenetic, metabolic states, proliferation, and DNA damage status). MIBI is currently limited to a ~50-antibody targets due to the difficulties in obtaining certainly isotopically-enriched compounds. If an increase in the number of parameters is required in future analyses, the EpicTag barcoding strategy could be adapted to alternative multiplex imaging technologies such as proteinbased CODEX (Goltsev et al. 2018), t-CyCIF (Lin et al. 2018), or Immuno-SABER (Saka et al. 2019), or nucleic acid-based FISSEQ (Lee et al. 2014), SeqFISH+ (Eng et al. 2019), MERFISH (Chen et al. 2015), SABER (Kishi et al. 2019), or Slide-seq (Rodriques et al. 2019). MIBI is applicable to 2D tissue sections, limiting preservation of relevant clonal cell spatial interactions in the z-axis such as irregularly-shaped tumor patches and vascular growth. Reconstruction based on serial sectioning might be an option (Catena et al. 2020) but remains time consuming. An appealing future direction will be leveraging on technologies for 3D multiplex imaging such as STARmap (Wang et al. 2018) or clearing-enhanced 3D microscopy (Li, Germain, and Gerner 2017).

The debarcoding pipeline also has limitations. Our strategy is based on integrating two independent initial steps: cell-based and pixel-based barcode assignment. Both provide valuable information. Cell-based assignment yields singlecell barcode classification but is prone to artifacts derived from the integration of signals on imperfectly segmented regions. Pixel-based assignment is not constrained by segmentation limitations and is thus a more faithful representation of barcode expression, but it does not provide single-cell barcode information and might be prone to artifacts in certain locations within the tissue. Integrating cell- and pixel-based barcode assignment provides robust classification for cells expressing proper barcode amounts, but classification of cells in noisy areas remains challenging. In addition, signal spillover from neighboring cells may be a confounding factor. Machine learning techniques (Moen et al. 2019) may enable more reliable debarcoding in noisy regions and an overall more efficient image processing.

In summary, the ability to track barcoded cancer cells *in situ*, together with the characterization of their spatial surroundings, opens new avenues for understanding tumor evolution. Barcoded cells can additionally be genetically modified for functional assessment of gene knockout, knockdown, overexpression, or mutation. Leveraging the capability to trace complex populations with distinct sets of genetic modifications, future work will be able to identify causal relations of genotypes and phenotypes to cellular neighborhoods. Lineage tracing is a blooming technology uniquely positioned to reveal the spatiotemporal dynamics of tissues at the singlecell scale (reviewed in (Wagner and Klein 2020)). EpicMIBI and other similar technologies (Frieda et al. 2017; Askary et al. 2020; Chow et al. 2021) will reveal principles and mechanisms of tissue developmental processes, in both normal and malignant growth, that will thus accelerate therapeutic discoveries.

## ACKNOWLEDGEMENTS

We thank Drs. Maria Angulo-Ibanez and Pablo Domizi for critical discussions and reading the manuscript, and all the members of the Bassik, Sage and Nolan labs for their help throughout this study. We thank Pauline Chu from the Stanford Histology Service Center, Catherine Carswell-Crumpton and Cheng Pan from the FACS Core of the Institute for Stem Cell Biology and Regenerative Medicine, Stanford’s animal mouse facility staff, Alyssa Ray and Nicole Wang for her help with administration and with figures, Thuyen Nguyen for her help with mice, and Angelica Trejo for technical contribution. Research reported in this publication was supported by the EMBO postdoctoral fellowship (ALTF 300-2017; to X.R.C), the Tobacco-Related Disease Research Program (T30FT0824; to A.P.D), the Stanford Dean’s Fellowship and the Leukemia & Lymphoma Society Career Development Program (to S.J.); grants of the National Institute of Health CA217450, and CA231997 (to J.S.), and 2U19AI057229-16, 5P01HL10879707, 5R01GM10983604, 5R33CA18365403, 5U01AI101984-07, 5UH2AR06767604, 5R01CA19665703, 5U54CA20997103, 5F99CA212231-02, 1F32CA233203-01, 5U01AI140498-02, 1U54HG010426-01, 5U19AI100627-07, 1R01HL120724-01A1, R33CA183692, R01HL128173-04, 5P01AI131374-02, 5UG3DK114937-02, 1U19AI135976-01, IDIQ17X149, 1U2CCA233238-01,1U2CCA233195-01, Task Order No. HHSN261100039 under Contract No. HHSN261201500003I (to G.P.N); grants of the Department of Defense W81XWH-14-1-0180, and W81XWH-12-1-0591 (to G.P.N.); grants of the Food and Drug Administration HHSF223201610018C, and DSTL/AGR/00980/01 (to G.P.N.); grant of the Cancer Research UK C27165/A29073 (to G.P.N.); grant of the Bill and Melinda Gates Foundation INV-002704 (to G.P.N.); grant of the Cancer Research Institute (to G.P.N.); grant of the Parker Institute for Cancer Immunotherapy (to G.P.N.); grant of the Kenneth Rainin Foundation 2018-575 (to G.P.N.); Celgene, Inc. 133826, and 134073 (to G.P.N.); the Rachford & Carlotta A. Harris Endowed Chair (to G.P.N.). This article reflects the views of the authors and should not be construed as representing the views or policies of the FDA, Department of Health and Human Services, BMGF or other institutions who provided funding.

## AUTHOR CONTRIBUTIONS

Conceptualization: X.R.C, A.P.D., J.S., G.P.N.
Methodology: A.P.D., X.R.C.
Software: S.J., X.R.C, A.P.D.
Formal Analysis: X.R.C., A.P.D, S.J., Y.B.
Investigation: A.P.D, X.R.C., S.J., Y.B., M.G., B.Z., M.M., G.C.
Writing – Original Draft: X.R.C., A.P.D, J.S., G.P.N.
Writing – Reviewing and Editing: all authors
Visualization: X.R.C, A.P.D., J.S., G.P.N.
Supervision: M.C.B., J.S., G.P.N.
Project Administration: J.S., G.P.N.
Funding Acquisition: J.S., G.P.N.

## MATERIALS AVAILABILITY

Plasmids generated in this study have been deposited to Addgene (Deposit 78757).

## DATA AND CODE AVAILABILITY

Original/source data and code for debarcoding can be accessed at https://data.mendeley.com/datasets/4y3fzctgxn/draft?a=042a495e-998a-4caf-b967-4030b7f5e7bf. After peer-review, the code for debarcoding generated during this study will be available on GitHub. After peer-review, original/source data for datasets 1 and 2 in the paper will be available on the public instance of MIBITracker at https://mibi-share.ionpath.com/.

## Materials & Methods

### Cell line model

NCI-H82 cells were purchased from ATCC (HTB-175™). Cells were cultured in RPMI media: RPMI 1640 (Corning, #15-040-CV) with 10% bovine growth serum (Thermo Fischer Scientific, #SH3054103HI), plus 1x penicillin/streptomycin plus glutamine (Thermo Fisher Scientific, #10378016). Cells tested negative for mycoplasma.

### Xenograft mouse model

Experiments with mice were conducted in accordance with Stanford University Animal Care and Use Committee guidelines. Cells were prepared in antibiotic-free media and mixed with Matrigel (BD biosciences, # 356237) at a 1:1 ratio. Aliquots of 200 μL (2-5 × 10^6^ cells) were injected subcutaneously into the flanks of NOD-scid IL2Rgamma^*null*^ mice (NSG mice). Tumors were collected once they reached a size of ~1 cm^3^.

### Vectors

EpicVectors are lentiviral vectors that express a puromycin resistance gene, T2A, and GFP or mCherry linked to a combination of three of six possible epitopes (VSVg, AU1, FLAG, StrepII, Prot C, HA). The vectors are deposited to Addgene (#162079-162118), and information available is in the Key Resources table.

### Lentiviral production

One day prior to transfection, 293T cells (ATCC, CRL-3216™) were seeded in DMEM (Thermo Fisher Scientific, #11965092) and 10% fetal bovine serum (Thermo Fisher Scientific, # 16000044) at 1 million cells per well of a 6-well plate. The following day, 293T cells were transfected with a mixture of 0.5 μg of each lentiviral EpicVector (Key Resources), 5 μg of 3^*rd*^ generation lentiviral vectors (at a 1:1:1 ratio of VSV-g, pMDLg/pRRE, and pRSV-Rev), 5 μL PEI (Polysciences, #23966-1) in 50 μL Opti-MEM (Thermo Fisher Scientific, # 31985088). The mixture was incubated 10 min and added dropwise to cells. The next day, the media was removed, and 2 mL of fresh antibiotic-free media was added to the cells. After 2 days, all viruses were collected, and cells were infected with 2 mL of virus per 4 million cells for each EpicVector, with the addition of polybrene (Millipore-Sigma, #TR-1003-G) at 8 μg/mL, bringing the total volume of media to 5 mL. The next day, the virus was removed, and fresh media containing penicillin/streptomycin (Thermo Fisher Scientific, #10378016) was added. After 2 days, multiplicity of infection was measured by FACS for GFP or mCherry. Puromycin was added for approximately 1 week until over 98% of cells were positive for GFP or mCherry.

### Ribonucleoprotein nucleofection

Three sgRNAs were designed to hybridize approximately 150 bases apart on the target of interest (see Key Resources) and were synthesized by Synthego. The three sgRNAs were resuspended in Tris-EDTA and were mixed at a 1:1:1 ratio. First, 12 μL of SE buffer (Lonza, #V4XC-1032) was added to each well of a 96-well v-bottom plate. Then 3 μL of sgRNA (300 pmol for all three sgRNAs) was added to the SE buffer. An aliquot of 0.5 μL of Alt-R® S.p. Cas9 (Integrated DNA Technologies, # 1081059) was added to 10 μL of SE buffer. Next the Cas9 was added to the sgRNA solution, mixed thoroughly, and incubated at 37 °C for 15 min to form the ribonucleoproteins (RNPs). NCI-H82 cells were pelleted, counted, and resuspended to 1 million cells per reaction in 5 μL of SE Buffer. Cells and the RNP solution were added to each well. Cells were immediately nucleofected using the Lonza 4D-Nucleofector*™* × Unit (Lonza, #AAF-1002X) with the EN150 program. After nucleofection, warm RPMI media was added to the cells. Cells were incubated at 37 °C for 15 min and then transferred to a 24-well plate. Editing efficiency was evaluated 4 days later by FACS or sequencing.

### Flow cytometry and cell sorting

Flow cytometry was conducted on a CytoFLEX (Beckman Coulter) and on a FACSAria (BD Biosciences). Analysis of the data was done using Cytobank and R analysis software.

### Antibody conjugation to isotopes

The antibodies listed in the Key Resources Table were conjugated to isotope-chelated Maxpar X8 polymers (Fluidigm, #201300) as previously described (G. Han et al. 2018). Briefly, 100 g of antibody in carrier-free PBS were partially reduced by TCEP treatment (Thermo Fisher Scientific, #77720) for 30 min at 37 °C. Reduced-antibodies were mixed with isotope-chelated polymers for 1.5 h at 37 °C. Antibody concentration was quantified via Nanodrop. Isotope-conjugated antibodies were diluted to >0.2 mg/mL in PBS antibody stabilizer (Thermo Fisher Scientific, #NC0414486) and stored at 4 °C.

### Palladium barcoding for CyTOF

Cells were fixed in freshly prepared 1.6% Paraformaldehyde (PFA) in 1X PBS (Thermo Fisher Scientific, #28906) for 20 min at room temperature. Cells were washed twice with ice-cold 1X PBS and once with ice-cold 0.02% Saponin (Millipore-Sigma, #S7900-100G) in 1X PBS for 20 min at 4 °C. Each palladium barcode (Fluidigm, #201060) was resuspended in 1 mL ice-cold 0.02% Saponin in 1X PBS and transferred to the selected sample. Cells were pipetted to ensure proper mixing with the palladium barcoding reagent and incubated for 15 min at room temperature. Cells were then washed twice with cell staining media (CSM; 0.5% bovine serum albumin (Millipore-Sigma, #A3059) and 0.02% sodium azide (Millipore-Sigma, #71289) in 1X PBS), and mixed into a single cell suspension.

### Cell preparation for CyTOF

Cells were fixed in freshly prepared 1.6% PFA in 1X for 20 min at room temperature. Cells were washed thrice with cold 1X PBS and permeabilized in pre-chilled 100% methanol for 20 min at 4 °C. Cells were washed once with 1X PBS and twice with CSM. Barcoded cells were stained for 3 h at 4 °C in CSM with the following cocktail of isotope-conjugated antibodies at 2 g/mL: anti-GFP, anti-AU1, anti-FLAG, anti-HA, anti-StrepII, anti-Prot C, anti-VSVg, and anti-cleaved caspase 3 (clone C92-605, BD Biosciences, #559565). After staining, cells were washed twice with CSM, once with 1X PBS, and incubated with 1X PBS containing 1.6% PFA and 1 M iridium-based DNA intercalator (Fluidigm, #201192B) for 16 h at 4 °C. After intercalation, cells were washed once with 1X PBS and thrice with distilled water before analysis. Data were collected with a CyTOF II mass cytometer (Fluidigm). The raw FCS files processing was performed in CellEngine (CellCarta) by gating out doublets based on cell length, debris based on iridium staining, and dead cell based on cleaved caspase staining. The resulting data were plotted using UMAP (*umap* R package v.0.2.7.0) (McInnes, Healy, and Melville 2018) and visually inspected to identify barcodes.

### Tumor preparation for CyTOF

Tumors were collected from the flanks of the mice and were minced using a razor blade. The samples were then placed to a 50 mL conical tube containing 9 mL of L15 media (Sigma Aldrich, # L1518) plus 1 mL enzyme mix (10 mL L15 with 85 mg Collagenase I (Millipore-Sigma, # C0130), 28 mg Collagenase II (Millipore-Sigma, # 6885), 85 mg Collagenase IV (Millipore-Sigma, # 5138), 12.5 mg Elastase (CellSystems, # LS002292), 12.5 mg DNAseI (Millipore-Sigma, #10104159) and filtered with a 0.22 m filter (Millipore-Sigma, # SLMP025SS)). The samples were incubated at 37 °C for 15 min in a slant with rotation. Then the mixture was filtered using a 70 m strainer (Fisher Scientific, # 08-771-2) into a new 50 mL tube. The samples were centrifuged at 400 xg for 5 min at room temperature. The supernatant was discarded, and 1 mL of red blood cell lysis buffer (Thermo Fisher Scientific, Cat# 00-4333-57) was added for 30 seconds. Then, 30 mL of 1XPBS were added and the samples were centrifuged at 400 g for 5 min at room temperature. Cells were collected in RPMI media without antibiotics supplemented with 10% DMSO, and frozen down to −80 °C until use.

### Cell pellet preparation for MIBI

All 20 barcoded NCI-H82 cell lines were mixed in a 15 mL conical tube and washed twice in cold 1X PBS by centrifuging at 125 xg for 5 min at 4 °C. The supernatant was decanted, 10 mL of freshly prepared 4% PFA in 1X PBS was carefully placed on top of the pellet, and the tube was placed on a rocker set at low speed for 16 h at 4 °C. The PFA-fixed cell pellet was pressed with a tip to cast it loose from the bottom of the tube and rinsed thrice by pouring and decanting 10 mL 1X PBS. The pellet was briefly dried on top of an absorbent piece of paper, transferred to a new 15 mL conical tube, submerged in pre-heated Histogel (Thermo Fisher Scientific, #NC9150318), and incubated for 10 min at 4 °C. The Histogel-embedded pellet was placed in a 9-spaces chamber, submerged in 80% ethanol, and FFPE processed.

### Tumor preparation for MIBI

Tumors were collected immediately after euthanasia, submerged in 10 mL of freshly prepared 4% PFA in 1X PBS (Thermo Fisher Scientific, #28906), and placed on a rocker set at low speed for 24 h at room temperature. Tumors were then rinsed thrice with 1X PBS, transferred to 80% ethanol, and FFPE processed.

### Gold-coated glass slide preparation

Superfrost Plus glass slides of 25 mm width and 75 mm length (Thermo Fisher Scientific, #12-550-15) were soaked in dish detergent, rinsed twice with distilled water, soaked in acetone, and air-dried with compressed air in a fume hood. Clean slides were coated with a 30 nm tantalum layer followed by a 100 nm gold layer. Coating was prepared at the Stanford Nano Shared Facility (Stanford, CA) and New Wave Thin Films (Newark, CA) as previously described (Ji et al. 2020; Keren et al. 2019).

### Vectabond treatment of gold-coated slides

Gold-coated slides were silanized by Vectabond treatment (Vector labs, #SP-1800-7). Using glass beakers, gold-coated slides were submerged in acetone for 5 min, placed in freshly prepared Vectabond solution (3.5 mL Vectabond and 175 mL acetone) for 30 min, and air-dried with compressed air in a fume hood. Slides were baked at 70 °C for 30 min and stored at room temperature.

### Sample preparation for MIBI

FFPE blocks were sectioned using a microtome into 5 m thin sections and placed on vectabond-treated gold-coated slides. Tissue sections were baked for 20 min at 70 °C, and immediately deparaffinized and rehydrated with fresh reagents as follows: xylene (x3), 100% ethanol (x2), 95% ethanol (x2), 80% ethanol, 70% ethanol, distilled water (x3). Each wash was performed for 3 min at room temperature with repetitive dipping using a linear stainer (Leica, #ST4020). Slides were transferred for heat-induced epitope retrieval to a PT module (Thermo Fisher Scientific, #A80400012) preheated to 75 °C. Samples in 1X Dako Target Retrieval Solution, pH 9 (Agilent, #S2375) were heated to 97 °C, stayed at the same temperature for 10 min, and then cooled down to 65 °C.

Slides were washed twice with 1X TBS IHC wash buffer with Tween 20 (Cell Marque, #935B-09) and 0.1% BSA (Thermo Fisher Scientific, #BP1600-100) for 5 min at room temperature. A hydrophobic barrier around the tissue was drawn using a PAP pen (Vector Laboratories, #H-4000). Tissue sections were blocked in 1X TBS IHC wash buffer with Tween 20, 2% normal donkey serum (Millipore-Sigma, #D9663-10ML) (MIBI wash buffer), 0.1% Triton X-100 (Millipore-Sigma, #T8787-100ML), and 0.05% sodium azide (Millipore-Sigma, #71289-50G) for 1 h at room temperature. Tissue sections were then stained for 16 h at 4 °C in 1X TBS IHC wash buffer with Tween 20, 3% normal donkey serum, and 0.05% sodium azide with the following cocktail of isotope-conjugated antibodies: anti-GFP, anti-AU1, anti-FLAG, anti-HA, anti-StrepII, anti-Prot C, anti-VSVg, anti-Vimentin, anti-HH3, anti-phosphoS28 HH3, anti-human-specific mitochondrial marker, anti-Ki67, anti-aSMA, anti-CD31, anti-Citrate synthase, anti-GLUT1, anti-phosphoS139 H2AX, anti-synaptophysin, anti-H3K27ac, anti-H3K4me2, and anti-H4K8ac.

After staining, slides were washed thrice with MIBI wash buffer for 5 min at room temperature, post-fixed with a solution of 2% glutaraldehyde (Electron Microscopy Sciences, #16120) and 4% PFA in 1X PBS for 5 min at room temperature and quenched with 100mM Tris pH 7.5 for 5 min at room temperature. Tissue sections were then dehydrated with fresh reagents as follows: 100mM Tris pH 7.5 (x2), distilled water (x3), 70% ethanol, 80% ethanol, 95% ethanol (x2), 100% ethanol (x2). Each wash was performed for 3 min at room temperature with repetitive dipping using a linear stainer. The slides were air-dried and stored at room temperature under vacuum until MIBI acquisition.

### MIBI data acquisition

MIBI data were acquired with a MIBI-TOF mass spectrometer (Ionpath) using a duoplasmatron ion source running research grade oxygen (Airgas, OX R80). We acquired two datasets for this study. Dataset 1 consists of the 16 tiles used for figures 3 to Dataset 2 consists of 14 tiles, 4 “Control” and 10 “PTEN”, used for Figure 6. Each tile acquired consisted of 9 FOV (Figure S3B). The acquisition parameters per FOV were the following:

1. Pixel dwell time: 7 ms
2. Image area: 400 m × 400 m
3. Image size: 512 × 512 pixels
4. Probe size: ~ 400 nm
5. Primary ion current: ~ 3.5 nA
6. Number of depths: 1

### Initial processing of MIBI images

Raw MIBI data were processed using MIBIAnalysis tools (https://github.com/lkeren/MIBIAnalysis) as previously described (Baranski et al. 2021; Keren et al. 2018). The mass spectra were calibrated using sodium and gold. The number of nearest neighborhoods for denoising was 25.

### Cell segmentation of MIBI images

MIBI image cell segmentation was obtained with Mesmer, a deep learning algorithm based on the DeepCell library (deepcell-tf 0.6.0) (Van Valen et al. 2016; Greenwald et al. 2021). The neural network weights for prediction were imported from https://deepcell-data.s3-us-west-1.amazonaws.com/model-weights/Multiplex_Segmentation_20200908_2_head.h5. Segmentation was computed using denoised and capped at the 99.7^*th*^ percentile images of HH3 and human-specific mitochondrial marker as input, to account for the nucleus and cytoplasm, respectively. Model_mpp in the python script multi-plex_segmentation.py was 1.8 for dataset 1 and 1.6 for dataset 2.

### Cell type annotation

Cell segmentation was used to extract cell counts per marker. Counts were normalized by cell size, scaled based on the median HH3 expression per cell in each tile, and transformed using an inverse hyperbolic sine (asinh) with cofactor of 0.05 (to account for the adjustment based on HH3 scaling).

Unsupervised single-cell clustering was performed using the FlowSOM R package (Van Gassen et al. 2015). The channels used for clustering were vimentin, phosphoS28 HH3, human-specific mitochondrial marker, Ki67, aSMA, CD31, citrate synthase, GLUT1, phosphoS139 H2AX, synaptophysin, H3K27ac, H3K4me2, and H4K8ac. Phenotypic SOM clusters were manually annotated based on visual inspection of the heatmap of marker expression per SOM cluster. Multiplex MIBI images were carefully compared to the SOM cluster maps, generated by coloring the cells with the respective SOM cluster, to assess accuracy and specificity of the clustering result and annotation. Single-cells were plotted using the umap R package (n_neighbors = 15 and min_dist = 0.01) and colored by SOM cluster to orthogonally inspect the clustering results and annotation. Single cells in properly annotated clusters were classified, and the remainder clusters were subjected to additional rounds of clustering until all cells were annotated. A post-clustering identification of endothelial cells was performed by selecting cells with low human-specific mitochondrial marker, small cell size, and high CD31 expression.

### CN analysis

Cell centroids were computed and the 30 nearest neighbors were selected for each cell. The frequencies of each SOM cluster were calculated per cell. Data from all tiles for each dataset were combined, and CNs were selected using k-means (k = 20) and labeled based on visual inspection of the heatmap of SOM cluster frequencies per CN. The CN maps were generated by coloring the cells with the respective CN.

### Debarcoding pipeline

Epitope debarcoding was performed by merging cell-based and pixel-based barcode assignment. The input for cell-based assignment of barcodes were the segmentation map and the six epitope MIBI images. Epitope counts within each segmented cell were extracted, normalized, and ordered to provide a barcode (1 to 20) to each cell.

The input for pixel-based assignment of barcodes were the six epitope MIBI images. A sliding window scan was applied to extract the counts of each epitope by centering a n × n window to each pixel. The size of the sliding window was an odd number (e.g., 3 × 3, 5 × 5, etc.). After applying a sliding window, the value of each epitope was capped at the 95^*th*^ percentile, and a barcode (1 to 20) was provided based on the 3 most expressed epitopes. A selection step of the third minus the fourth most expressed epitopes being higher than 0.05 was applied to account for blank pixels. The data were plotted back to the 2D tissue space and groups of pixels larger than 30 and sharing a barcode were selected. The unselected pixels were subjected to extra rounds of scanning with sliding window of larger size, up to sliding windows of 11 × 11.

To merge cell- and pixel-based barcode assignments, each cell within the tile is classified in one of six categories.

1. Category a: a cell with the same cell-based barcode as its pixel-based barcode retains that barcode.
2. Category b: a cell with a cell-based barcode without pixel-based barcode is assigned the cell-based barcode.
3. Category c: a cell without a cell-based barcode without pixel-based barcode results in a cell without barcode.
4. Category d: a cell without a cell-based barcode that has 50% or more of its area within an area of the same pixel-based barcode is assigned the pixel-based barcode.
5. Category 3e: A cell with cell-based barcode within areas with two or more pixel-based barcodes results in a cell with the barcode of the area from which the cell shares the highest surface percentage.
6. Category f: A cell with cell-based barcode overlapping with an area presenting a distinct pixel-based barcode results in a cell keeping the cell-based barcode if the percentage of shared area is less than 50%. Alternatively, it results in a cell with pixel-based barcode if the percentage of shared area is 50% or more.

To identify subclonal tumor patches, a barcode based on the merging step was provided to pixels related to single cells, and a barcode from the pixel-based barcode assignment was provided to pixels unrelated to single cells. Single cells within groups of pixels sharing a barcode were provided with the same patch identification number.

**Statistical analysis**

Statistical analysis was conducted using R. Significance is calculated by ANOVA within and between groups and adjusted by Bonferroni. P-adjusted: 0.05-0.01:*, 0.01-0.001:**, <0.001:***.

### Data visualization

MIBI data visualization was performed in ImageJ or Ionpath MIBITracker. Plots were created using the ggplot2 R package (Wilkinson 2011). Figures 1A, 2A, 6A, and S2A were created in part using BioRender.com. All figures were prepared using Illustrator (Adobe).

---

**Figure S1.**
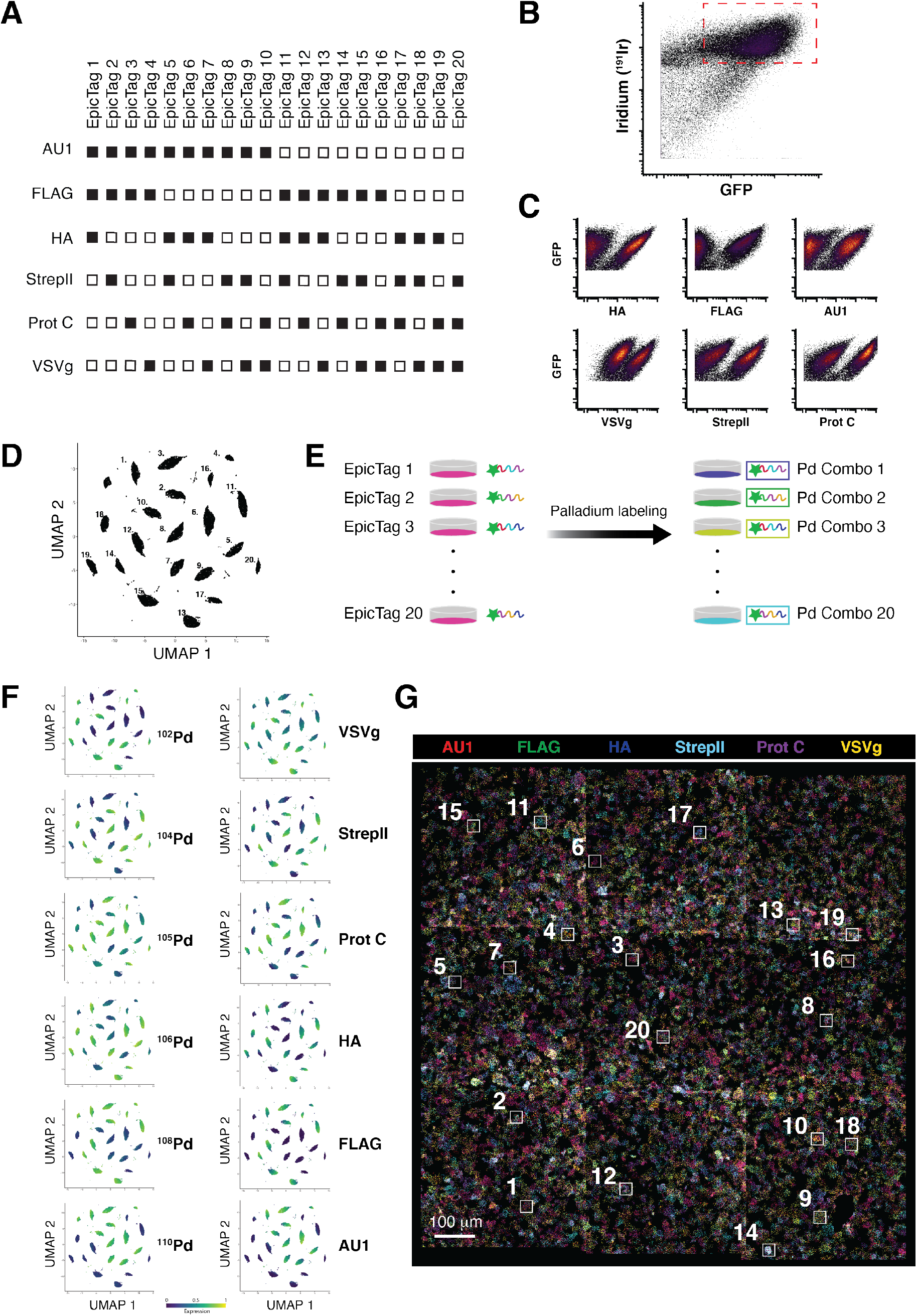
Validation of barcode expression in NCI-H82 SCLC cells, related to Figure 1. (**A**) Schematic of the expected epitopes (rows) for each combination (columns) in a 6-choose-3 strategy. A filled square indicates presence of the epitope in the combination. An empty square indicates absence of the epitope in the combination. (**B**) GFP and iridium (^191^ Ir)-based DNA intercalator were measured by mass cytometry in a pooled population of 20 barcoded NCI-H82 cell lines. The dashed red box indicates live cells expressing high levels of GFP that were selected for further analysis. (**C**) Biaxial plots showing positive and negative populations for each of the epitopes, validating epitope expression and antibody staining. GFP and epitopes were measured by mass cytometry in the pooled population of 20 barcoded NCI-H82 cell lines (n = 45,700). (**D**) UMAP of cells (n = 45,700) grouped by epitope expression (AU1, FLAG, HA, StrepII, Prot C, and VSVg). Each point represents a cell. Each number indicates a barcode combination (see epitope column in “F”). (**E**) Schematic of the palladium staining. Each of the 20 epitope combinations is labeled by one unique palladium combination. (**F**) UMAP of cells (n = 45,700) grouped by epitope expression and colored by epitope (AU1, FLAG, HA, StrepII, Prot C, and VSVg) or palladium isotope intensity. Pairs of palladium isotope and epitope in each row show similar patterns indicating a unique palladium and epitope combination for each of the 20 cell lines. (**G**) Representative overlay of MIBI images (AU1: red; FLAG: green; HA: blue; Strep II: cyan; Prot C: magenta; VSVg: yellow) of a cell pellet consisting of 50% wild-type NCI-H82 cells and 50% of a pooled population of 20 barcoded NCI-H82 cell lines. The white squares are enlarged in Figure 1E and their associated numbers indicate the barcode combination. The multicolor image in Figure 1D is an enlarged version of the bottom right of this image. Scale bars: 100 μm.

**Figure S2.**
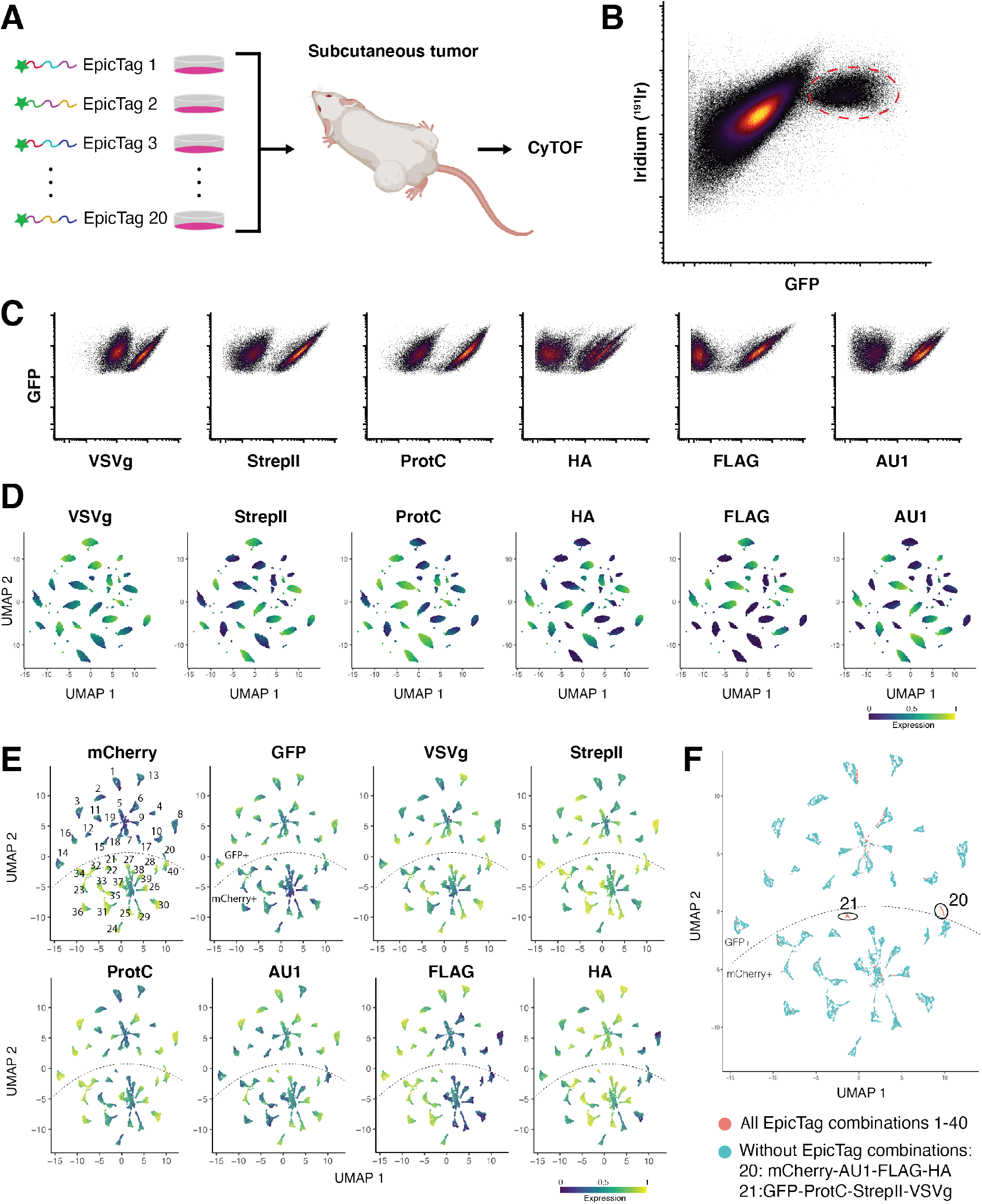
Validation of barcode expression in NCI-H82 SCLC xenografts, related to Figure 2. (**A**) Workflow for mass cytometry analysis of NCI-H82 xenografts. (1) Cell lines expressing unique combinations of epitopes are generated based on an 6-choose-3 strategy and grown *in vitro* in an independent flask. (2) Barcoded cells are mixed and injected subcutaneously into the flanks of a mouse. (3) Tumors are processed to generate a single-cell suspension, stained with an isotope-conjugated antibody cocktail, and analyzed by mass cytometry. (**B**) GFP and iridium (^191^ Ir)-based DNA intercalator measured by mass cytometry in a single-cell suspension from barcoded NCI-H82 tumors. The dashed red ellipse indicates live cells that express high levels of GFP that were selected for further analysis (Figures S2C and D; n = 39,630). (**C**) Biaxial plots showing positive and negative populations for each of the epitopes, validating epitope expression and antibody staining in cells from barcoded NCI-H82 tumors. GFP and epitopes were measured by mass cytometry in the single-cell suspensions from tumors. (**D**) UMAP of all cells from the dashed red ellipse in panel B (n = 39,630) colored by epitope expression (AU1, FLAG, HA, StrepII, Prot C, and VSVg). Each point represents a cell. Each epitope barcode is a unique group. (**E**) UMAP of cells from a pooled population of 40 barcoded NCI-H82 cell lines analyzed by mass cytometry grouped by protein and epitope (GFP, mCherry, AU1, FLAG, HA, StrepII, Prot C, and VSVg) and colored by their intensity. Each point represents a cell. Each number indicates a barcode combination. The dashed lines separate GFP-positive from mCherry-positive cells. (**F**) Overlay of 40 barcoded NCI-H82 cell lines (red) and a pooled population of 38 barcoded NCI-H82 cell lines (blue). The latter includes the same cell lines as were mixed in the 40-line pool except for the line expressing GFP with a tag consisting of Prot C, StrepII, and VSVg (combination 20) and the line expressing mCherry with a tag consisting of AU1, FLAG, and HA (combination 21). Each number indicates a barcode combination. The dashed lines separate GFP-positive from mCherry-positive cells.

**Figure S3.**
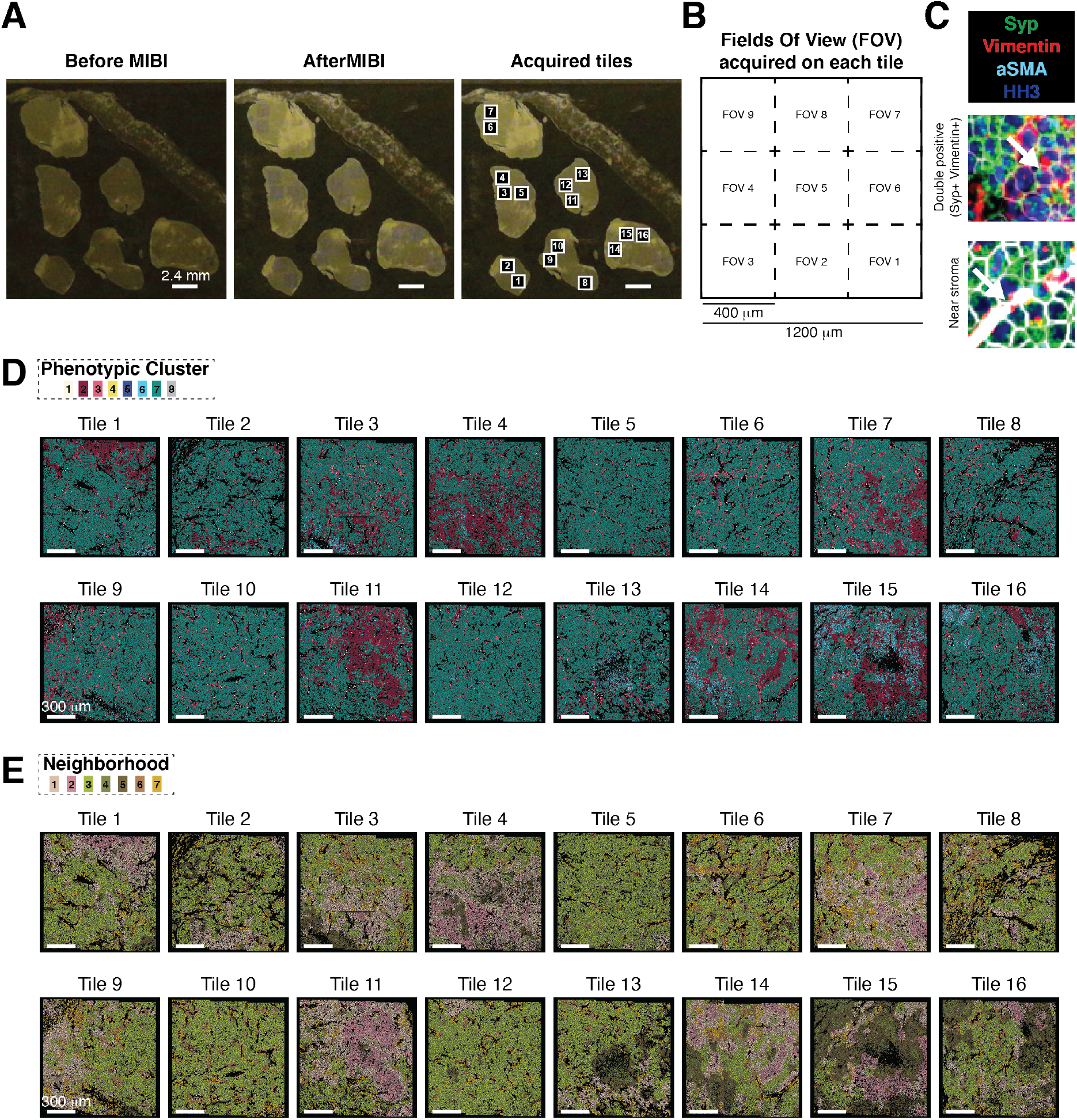
Overview of phenotypic clusters and CNs in NCI-H82 xenografts, related to Figure 3. (**A**) Left: Representative image of a barcoded NCI-H82 tumor cut and mounted onto a glass slide before MIBI. Middle: The same image after MIBI acquisition; darker regions indicate the areas acquired. Right: Numbered areas within each tumor indicate the acquired tiles. Data from this dataset are shown in Figures 3 to 5. Scale bar: 2.4 mm. (**B**) Schematic representation of tile acquisition. Each tile has a size of 1200 × 1200 ?m and is composed of nine squared FOVs of size 400 × 400 ?m. Each dashed square represents a FOV. The order of FOV acquisition is indicated by the numbers within each square. (**C**) Representative overlaid MIBI images of synaptophysin (SYP; green), vimentin (red), aSMA (cyan), and HH3 (blue) staining of NCI-H82 cells. Top image: Region of vimentin co-expression with synaptophysin (white arrow). Bottom image: Region of vimentin expression in proximity to stroma (white arrow). White lines are segmented cells. (**D**) Phenotypic cluster maps for each tile. Numbers in the legend refer to phenotypic clusters defined in Figure 3F. Tile 1 is shown in Figure 5A. Tile 5 and 15 are shown in Figure 3I. Scale bar: 300 μm. (**E**) Neighborhood maps for each tile on the dataset. Numbers in the legend refer to CNs defined in Figure 3K. Tile 1 is shown in Figure 5A. Tile 5 and 15 are shown in Figure 3L. Scale bar: 300 μm.

**Figure S4.**
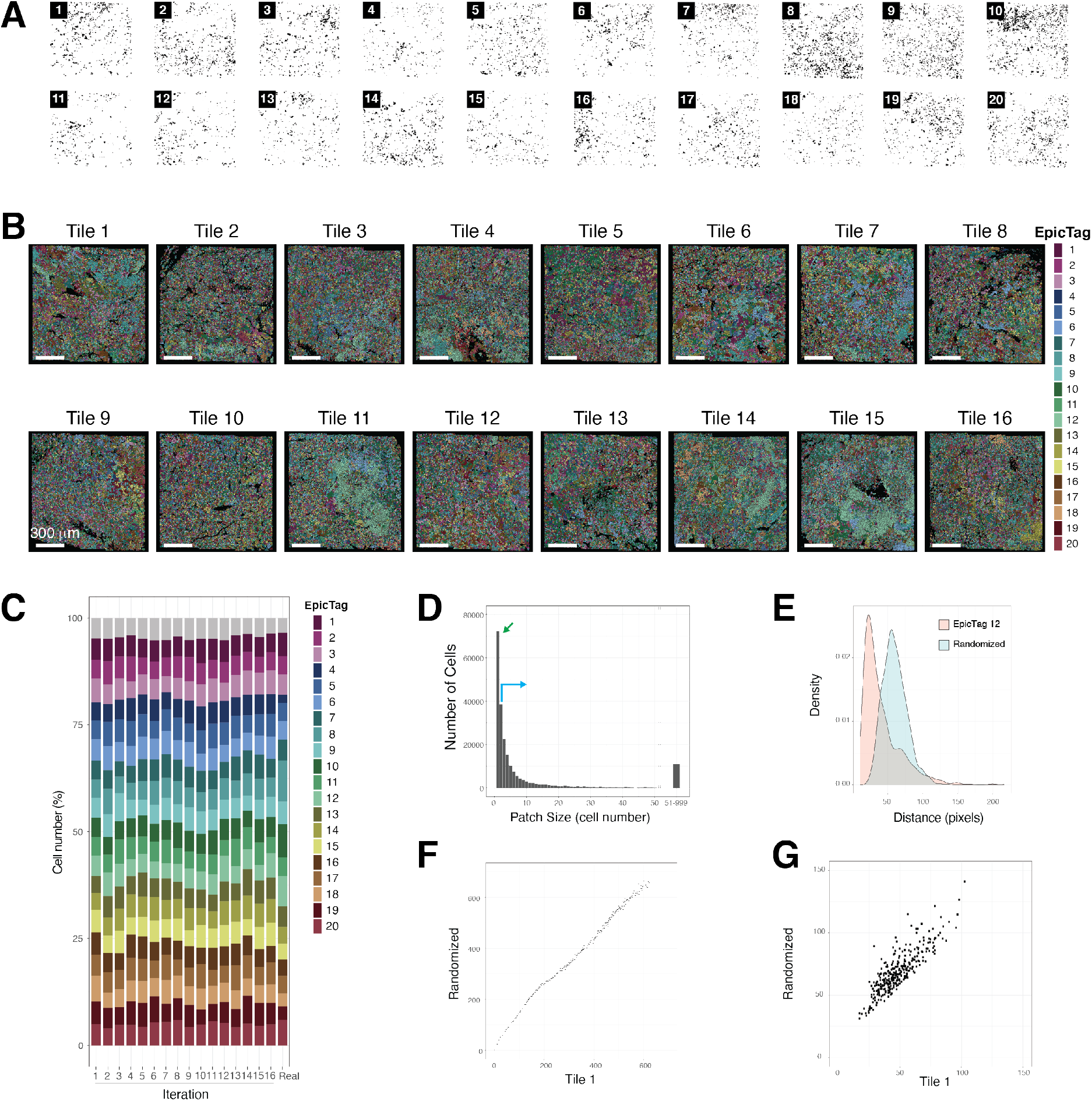
Overview of EpicTags in NCI-H82 xenografts, related to Figure 4. (**A**) Positive cells for each barcode in the tile shown in Figure 4B. The number within the black square indicates the barcode. (**B**) Subclonal tumor maps for each tile on the dataset. Numbers and colors in the legend refer to the barcodes described in Figure S1A. Tile 1 is shown in Figure 5A. Tile 14 is shown modified in Figure 5D. Scale bar: 300 μm. (**C**) Computational simulation of frequencies of each EpicTag barcode after a random initial seeding. (**D**) Bar graph showing the number of cells for each patch size in the dataset. The green arrow indicates individually distributed cells. The blue green indicates cells in patches. The x-axis is log10 transformed on the right. (**E**) Histogram showing the distances of each cell to its fifth nearest neighbor. The pink distribution represents the distribution of cells in the tumor for barcode 12 in tile 1 and the blue distribution represents a randomized sample in tile 1 (with the same number of cells of barcode 12). (**F**) Dot plot for barcode 12 in tile 1 showing the mean of the distances of each cell to its *k* nearest neighbor in the tumor and in randomized data. *k* ranges from 1 to 300. (**G**) Dot plot showing the mean of the distances of each cell to its *k* nearest neighbor in the tumor and in randomized data for all 20 barcodes in each of the 16 tiles (n = 320).

**Figure S5.**
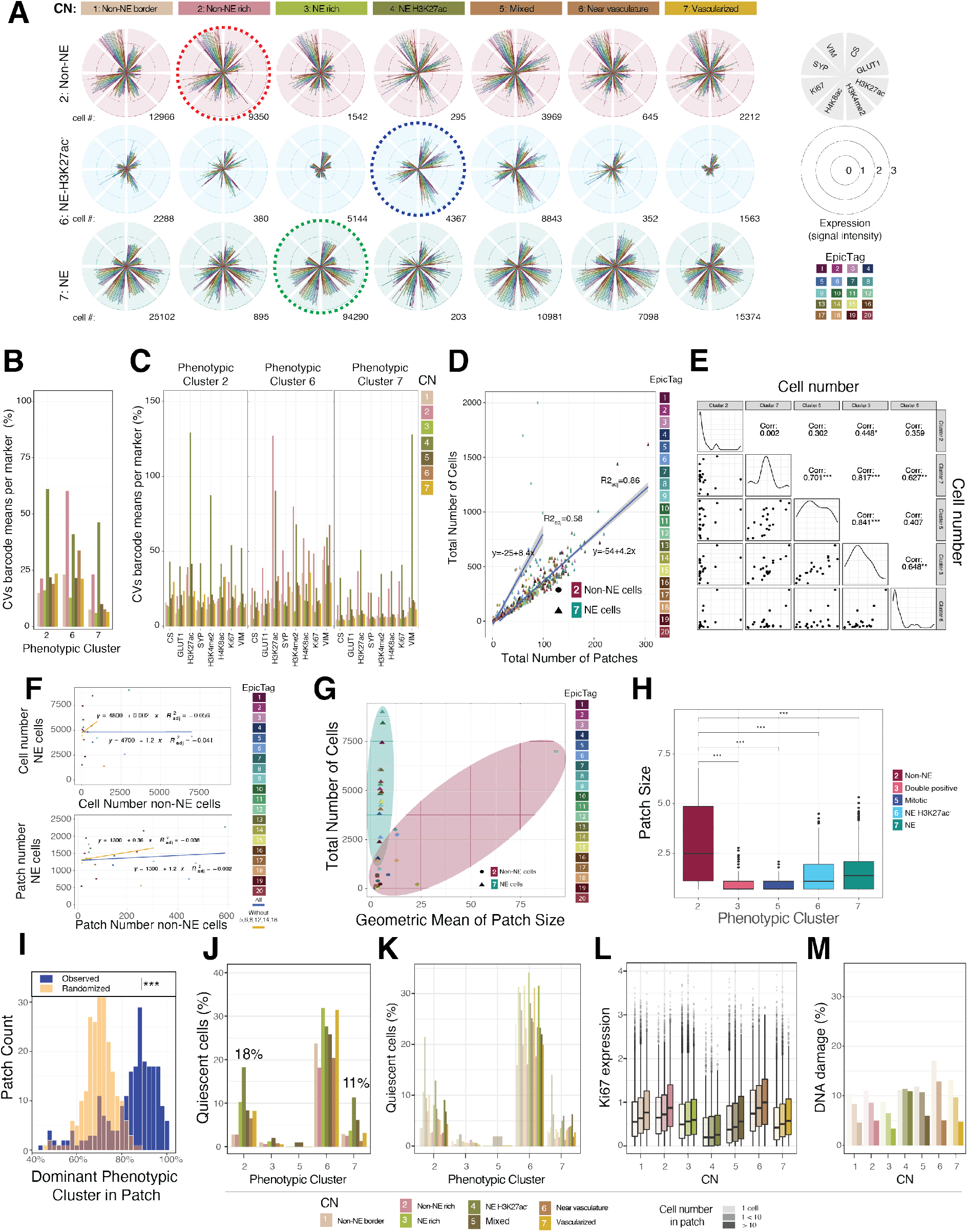
Functional analyses of phenotypic SOM clusters and CNs in NCI-H82 xenografts, related to Figure 5. (**A**) Polar plots illustrating marker expression (VIM, SYP, Ki67, H4K8ac, H3K4me2, H3K27ac, GLUT1, CS) across clusters and CNs. Concentric circles represent marker expression. Dashed lines indicate homotypic CNs (red: cluster 2 – CN 2, blue: cluster 6 – CN 4, green: cluster 7 – CN 3). (**B**) Average marker CV across clusters in CNs. The CV was calculated from means of expression in barcoded cells; the mean of all CVs is plotted. (**C**) CVs across clusters and CNs. (**D**) Representation of cell number and patches per tile. The total number of barcoded cells and patches per tile separated by clusters 2 (non-NE, non-neuroendocrine) and 7 (NE). Linear models fit the data for NE and non-NE cells with R^2^ *adj* = 86% and 58%, respectively. (**E**) Correlation of barcodes between NE and non-NE cells. Cell number (top) and patch number (bottom) of NE and non-NE cells are plotted per barcode. Linear models and correlations of the total data (blue) and data without combinations 5, 4, 8, 12, 14, and 18 (orange) are displayed. (**F**) Correlations, histograms, and points for each barcode for all clusters. (**G**) Geometric means of patches size across all barcodes for cluster 7 (NE) and cluster 2 (non-NE). (**H**) Patch sizes across all tiles per barcode for clusters 2, 3, 5, 6, and 7 (human cell clusters); statistical comparison was between cluster 2 and the remaining clusters. (**I**) Percentage of the dominant cluster within a patch (blue) and a randomized assignment of clusters within a patch (orange). (**J**) Quiescent cells per CN for each cluster. (**K**) Quiescent cells per cluster in each CN. Data were separated by patch size: 1 cell, 1 to 10 cells, and over 10 cells. (**L**) Cell proliferation as indicated by Ki67 expression for each cluster and CN. (**M**) Percentage of cells with DNA damage per cluster and CN.

**Figure S6.**
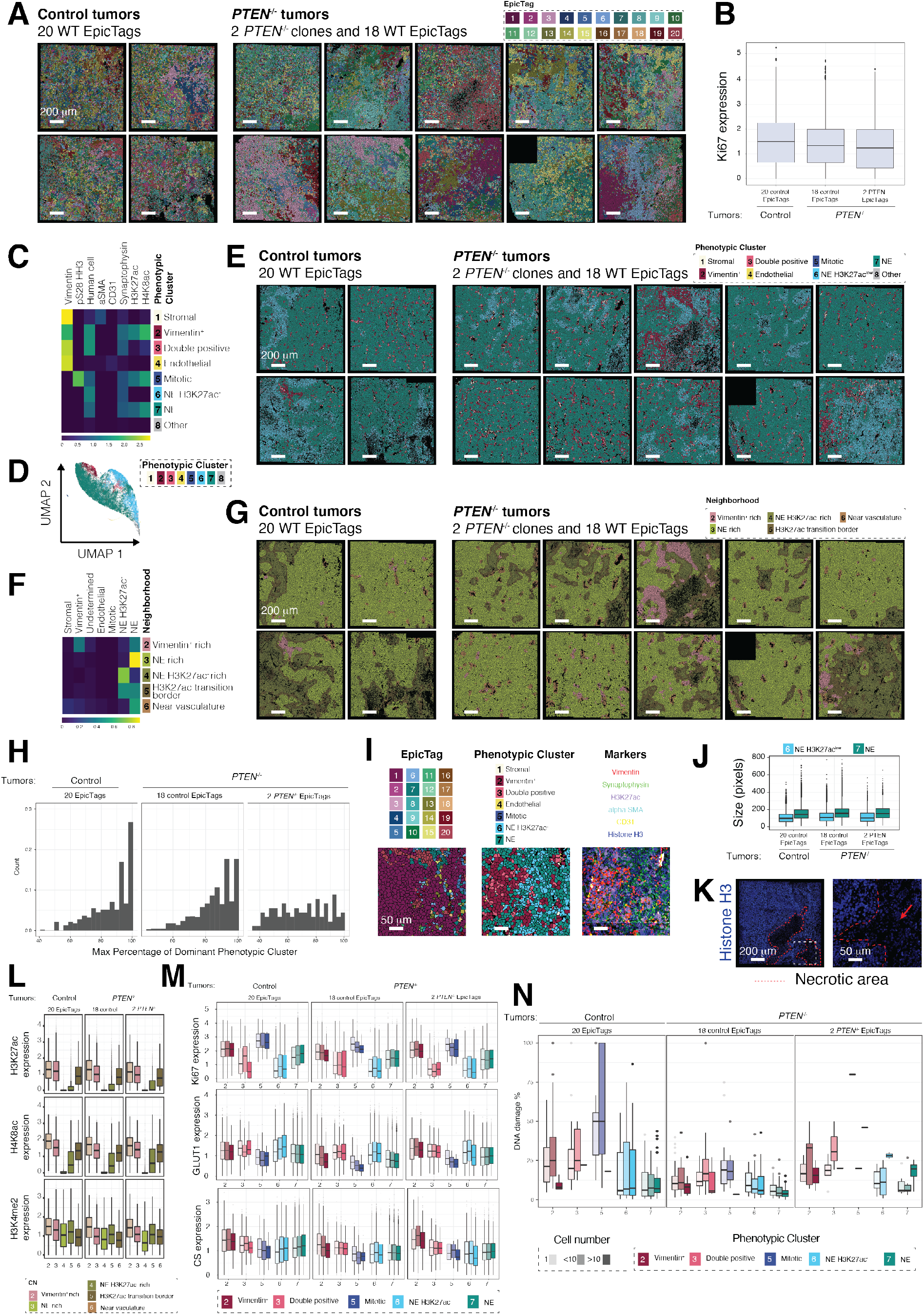
Functional analyses of control and PTEN^*−/−*^ tumors, related to Figure 6. (**A**) Four images from “Control” tumors consisting of 20 wild-type EpicTag barcoded cells, and ten images of “*PTEN^−/−^*” tumors consisting of *PTEN* knockout cells (EpicTags 1 and 20), unedited but nucleofected cells (EpicTags 2, 4, 14, 16, 17, and 18), and wild-type cells (EpicTags 3, 5, 6, 7, 8, 9, 10, 11, 12, 13, 15, and 19). The legend indicates the color for each EpicTag barcode. Numbers refer to the barcodes described in Figure S1A. All tiles are shown modified in Figure 6I. Scale bars: 200 μm. (**B**) Cell proliferation as indicated by Ki67 expression for cells in “Control” and “*PTEN^−/−^*” tumors. For the “*PTEN^−/−^*” tumors, proliferation of control cells (wild-type cells, and unedited but nucleofected cells) and *PTEN^−/−^* cells (“PTEN”) are shown. (**C**) A heatmap of the mean marker expression for the eight clusters (phenotypic cluster; rows) and the eight markers that define them (columns). The intensities of each marker are defined by the color bar at the bottom (normalized counts). (**D**) UMAP of all cells in the dataset (n = 194,821) colored by their phenotypic cluster. Each point represents a cell. Cells were grouped based on the expression of 13 markers (Vimentin, pS28 HH3, human cell marker, Ki67, aSMA, CD31, Citrate Synthase, GLUT1, pS139 H2AX, Synaptophysin, H3K27ac, H3K4me2, and H4K8ac). (**E**) Phenotypic cluster maps for tiles of the “Control” and “*PTEN^−/−^*” tumors. Numbers in the legend refer to phenotypic clusters defined in Figure S6C. Scale bars: 200 μm. (**F**) A heatmap of the frequencies of each phenotypic cluster (columns) for each of the seven CNs (rows). The normalized intensity of each marker is defined by the color bar. (**G**) Neighborhood maps for tiles of “Control” and “*PTEN^−/−^*” tumors. Numbers in the legend refer to CNs defined in Figure S6F. Scale bars: 200 μm. (**H**) Percentages of the dominant clusters within a patch for cells in “Control” and “*PTEN^−/−^*” tumors. For “*PTEN^−/−^*” tumors, percentages in control cells (wild-type cells, and unedited but nucleofected cells) and *PTEN^−/−^* cells (“PTEN”) are shown. (**I**) Subclonal tumor map, phenotypic cluster map, and a composite image of vimentin (red), synaptophysin (green), H3K27ac (violet), alpha SMA (cyan), CD31 (yellow), and Histone H3 (blue) in a representative region of a “PTEN” NCI-H82 xenograft (Tile 14, FOV 3 and 4). Bottom row images are enlarged representations of red dashed squares in the top row. White arrows indicate representative H3K27ac^−^, H4K8ac^+^, and H3K4me2^+^ cells enriched in CN 5. Scale bars: 50 μm. (**J**) Cell size by pixel count for NE and NE H3K27ac^*low*^ cells in “Control” and “*PTEN^−/−^*” tumors. For the “*PTEN^−/−^*” population, cell sizes of control cells (wild-type cells, and unedited but nucleofected cells) and *PTEN^−/−^* cells (“PTEN”) are shown. (**K**) Representative image of Histone H3 staining in a representative region of a “*PTEN^−/−^*” NCI-H82 xenograft (Tile 6). Image on the right is an enlarged representation of the white dashed square in the image on the left. Areas within the dashed red lines resemble necrotic areas. A red arrow indicates a region in the enlarged image that resembles a necrotic area. Scale bars: 200 μm (left) and 50 μm (right). (**L**) H3K27ac, H4K8ac, and H3K4me2 expression per CN in “Control” and “*PTEN^−/−^*” tumors. (**M**) Ki67, GLUT1, and citrate synthase (CS) expression per cluster in “Control” and “*PTEN^−/−^*” tumors. For the “*PTEN^−/−^*” population, expression is shown for control cells (wild-type cells, and unedited but nucleofected cells) and *PTEN^−/−^* cells (“PTEN”). (**N**) Percentage of cells with DNA damage per cluster in “Control” and “*PTEN^−/−^*” tumors. For the “*PTEN^−/−^*” population, percentages are given for control cells (wild-type cells, and unedited but nucleofected cells) and *PTEN^−/−^* cells (“PTEN”). For each cluster the data were separated by patch size: 1 cell, 1 to 10 cells, and over 10 cells.

